# Structural basis of transcription inhibition by fidaxomicin (lipiarmycin A3)

**DOI:** 10.1101/237123

**Authors:** Wei Lin, Kalyan Das, David Degen, Abhishek Mazumder, Diego Duchi, Dongye Wang, Yon W. Ebright, Richard Y. Ebright, Elena Sineva, Matthew Gigliotti, Aashish Srivastava, Sukhendu Mandal, Yi Jiang, Yu Liu, Ruiheng Yin, Zhening Zhang, Edward T. Eng, Dennis Thomas, Stefano Donadio, Haibo Zhang, Changsheng Zhang, Achillefs N. Kapanidis, Richard H. Ebright

**Affiliations:** Waksman Institute and Department of Chemistry, Rutgers University, Piscataway NJ 08854, USA; Rega Institute and Department of Microbiology and Immunology, KU Leuven, 3000 Leuven, Belgium; Department of Physics, University of Oxford, Oxford OX1 3PU, United Kingdom; The National Resource for Automated Molecular Microscopy, Simons Electron Microscopy Center, New York Structural Biology Center, New York NY 10027, USA; Center for Integrative Proteomics, Rutgers University, Piscataway NJ 08854, USA; NAICONS Srl., 20139 Milan, Italy; South China Sea Institute of Oceanology, Chinese Academy of Sciences, Guangzhou 510301, China

## Abstract

Fidaxomicin is an antibacterial drug in clinical use in treatment of *Clostridium difficile* diarrhea^1–2^. The active pharmaceutical ingredient of fidaxomicin, lipiarmycin A3 (Lpm)^1–4^, is a macrocyclic antibiotic with bactericidal activity against Gram-positive bacteria and efflux-deficient strains of Gram-negative bacteria^1–2, 5^. Lpm functions by inhibiting bacterial RNA polymerase (RNAP)^6–8^. Lpm exhibits no cross-resistance with the classic RNAP inhibitor rifampin (Rif)^7, 9^ and inhibits transcription initiation at an earlier step than Rif^8–11^, suggesting that the binding site and mechanism of Lpm differ from those of Rif. Efforts spanning a decade to obtain a crystal structure of RNAP in complex with Lpm have been unsuccessful. Here, we report a cryo-EM^12–13^ structure of *Mycobacterium tuberculosis* RNAP holoenzyme in complex with Lpm at 3.5 Å resolution. The structure shows that Lpm binds at the base of the RNAP “clamp,” interacting with the RNAP switch region and the RNAP RNA exit channel. The binding site on RNAP for Lpm does not overlap the binding sites for other RNAP inhibitors, accounting for the absence of cross-resistance of Lpm with other RNAP inhibitors. The structure exhibits an open conformation of the RNAP clamp, with the RNAP clamp swung outward by ~17° relative to its position in catalytically competent RNAP-promoter transcription initiation complexes, suggesting that Lpm traps an open-clamp conformational state. Single-molecule fluorescence resonance energy transfer^14^ experiments confirm that Lpm traps an open-clamp conformational state and define effects of Lpm on clamp opening and closing dynamics. We propose that Lpm inhibits transcription initiation by trapping an open-clamp conformational state, thereby preventing simultaneous engagement of transcription initiation factor σ regions 2 and 4 with promoter -10 and -35 elements. The results provide information essential to understanding the mode of action of Lpm, account for structure-activity relationships of known Lpm analogs, and suggest modifications to Lpm that could yield new, improved Lpm analogs.

---

We determined a structure of *Mycobacterium tuberculosis (Mtb)* RNAP σ^A^ holoenzyme in complex with Lpm by use of cryo-EM with single particle reconstruction (Fig. 1; Extended Data Figs. 1–3). Density maps showed unambiguous density for RNAP, including the taxon-specific, *Mycobacterium-specific* sequence insertion^15^ (β’MtbSI or “gate”), for σ conserved regions 2, 3, and 4 (σR2, σR3, and σR4), and for Lpm. The mean resolution of the structure is 3.5 Å (Extended Data Fig. 1e). Local resolution ranges from ~2.5-3.8 Å in central parts of the structure, including RNAP residues close to Lpm, to ~5-6.5 Å in peripheral parts of the structure, including *β’MtbSI* and σR4 (Fig. 1a; Extended Data Figs. 1f, 2a). Local B-factors range from ~0-100 Å^2^ in central regions to ~300-400 Å^2^ in peripheral regions (Extended Data Fig. 2b). Density maps show clear density for backbone and sidechain atoms of RNAP and individual functional groups of Lpm (Extended Data Fig. 2c-d).

**Figure 1.**
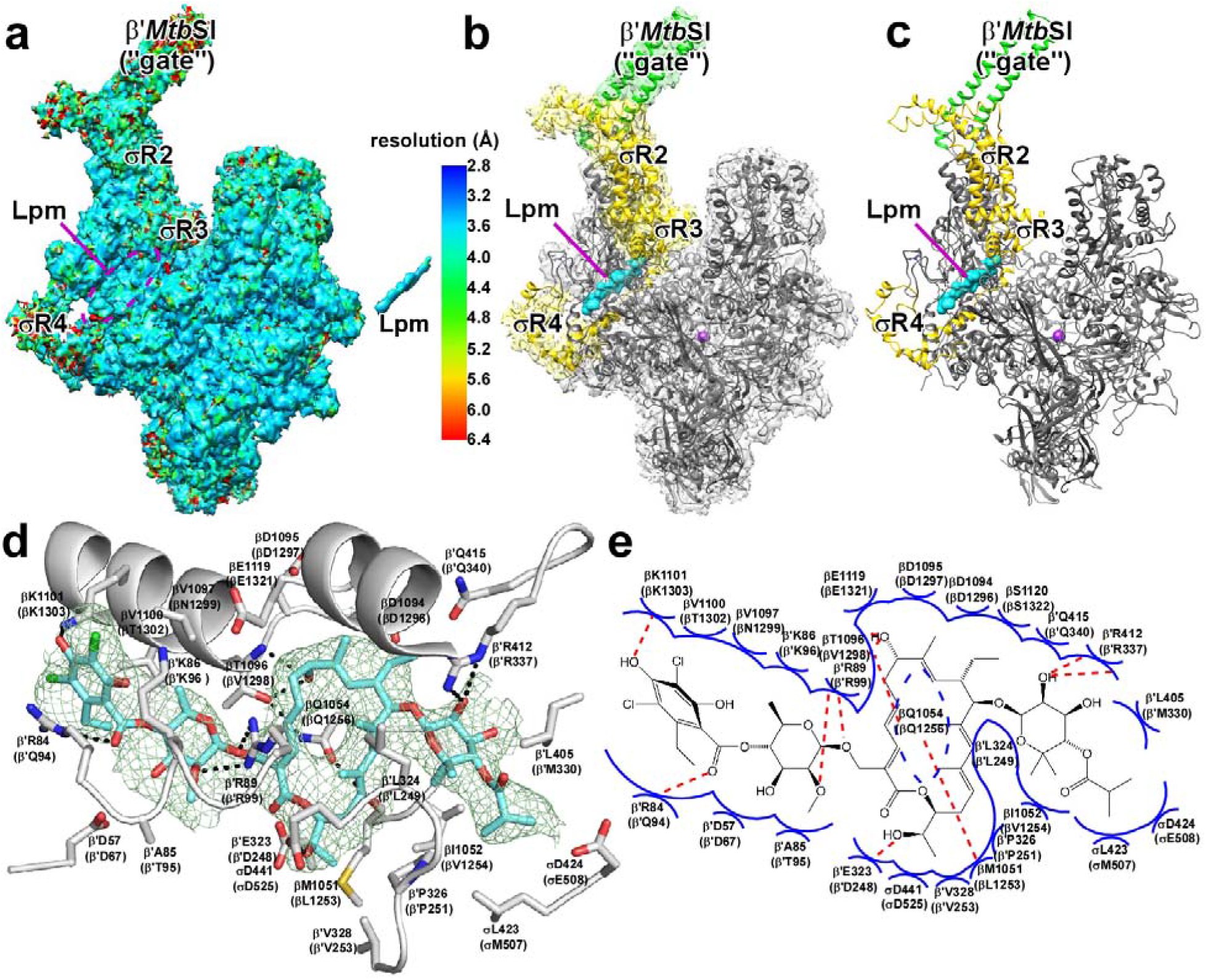
Structure of *Mtb* RNAP-Lpm. **a**, Density map for *Mtb* RNAP-Lpm (left) and Lpm (right), colored by local resolution. **b**, Density and atomic model for *Mtb* RNAP-Lpm. Gray, RNAP core other than β’MtbSI; green, β’MtbSI; yellow, σ; green mesh, density map for Lpm; cyan, Lpm; violet sphere, RNAP active-center Mg^2+^. **c**, Atomic model for *Mtb* RNAP-Lpm. Colors as in **b. d**, RNAP-Lpm interactions (residues numbered as in *Mtb* RNAP and, in parentheses, as in *Escherichia coli* RNAP). Gray ribbons, RNAP backbone; gray sticks, RNAP carbon atoms; cyan sticks, Lpm carbon atoms; red, blue, and green sticks, oxygen, nitrogen, and chlorine atoms; dashed lines, H-bonds. **e**, Summary of RNAP-Lpm interactions. Blue arcs, van der Waals interactions; red dashed lines, H-bonds.

The cryo-EM structure shows that the Lpm binding site is located at the base of the RNAP clamp^14, 16–17^ and encompasses RNAP clamp α-helices βa16α1 and β’a4α1 and RNAP switch-region switches SW2, SW3, and SW4^5, 17^ (Fig. 1; Extended Data Fig. 3). Lpm binds to RNAP in a fully-extended conformation similar to conformations in crystal structures of Lpm alone (Extended Data Fig. 3a). All five structural moieties of Lpm--isobutyryl, β-D-noviosyl, macrocycle, β-D-rhamnosyl, and homodichloro-orsellinyl--interact with RNAP (Fig. 1d-e; Extended Data Figs. 1a, 3b-c). Lpm makes H-bonds with six RNAP residues: βT1096 (V1298), βK1101 (K1303), β’R84 (Q94), β’R89 (R99), β’E323 (D248), and β’R412 (R337) (residues numbered as in*Mtb* RNAP and, in parentheses, as in *Escherichia coli* RNAP). Alanine substitutions of these RNAP residues results in Lpm-resistance, confirming the interactions are functionally relevant (Extended Data Fig. 3d).

Sequence alignments show that residues of RNAP that contact Lpm are conserved in Gram-positive and Gram-negative bacterial RNAP, but are not conserved in human RNAP I, II, and III (Extended Data Fig. 4a-b), accounting for the ability of Lpm to inhibit both Gram-positive and Gram-negative bacterial RNAP but not human RNAP I, II, and III (Extended Data Fig. 4c).

The cryo-EM structure accounts for structure-activity relationships of Lpm analogs produced by metabolism^18^, precursor feeding^19^, mutasynthesis^20–21^, and semi-synthesis (Fig. 1d-e; Extended Data Figs. 3b-c, 5). The structure accounts for structure-activity relationships indicating functional importance of the Lpm isobutyryl moiety (b1 vs. Lpm), β-D-noviosyl moiety (c1 vs. b2 and a6); macrocycle and C25 hydroxyl therein (a5 vs. Lpm), β-D-rhamnosyl moiety and C6’ methyl therein (e1 vs. d1 and a7; a6 vs. Lpm), and homodichloro-orsellinyl moiety and C8” methyl, C3” chlorine, and C5” chlorine therein (d1 vs. Lpm; a2-a4 vs. Lpm; a8 vs. a6). The observation that the Lpm C4” hydroxyl is exposed to solvent at the exterior of RNAP (Extended Data Fig. 3b-c) predicts that substituents--including large substituents--may be introduced at the C4” hydroxyl oxygen without eliminating ability to inhibit RNAP, provided the H-bond-acceptor character of the oxygen is maintained. Validating this prediction, we have prepared a novel Lpm analog having benzyl appended at the C4” hydroxyl oxygen by semi-synthesis from Lpm and have found the analog to retain high activity in inhibiting RNAP (Extended Data Fig. 5, analog a1).

The binding site on RNAP for Lpm (cyan in Fig. 2a) does not overlap the binding sites of other RNAP inhibitors, including rifampin and sorangicin^22–23^ (Rif and Sor); GE23077 and pseudoridimycin^24–25^ (GE and PUM), CBR703 and D-AAP-1^15, 26–27^ (CBR and AAP), salinamide^28^ (Sal), streptolydigin^29–30^ (Stl), and myxopyronin, corallopyronin, ripostatin, and squaramide^31–33^ (Myx, Cor, Rip, and SQ) (Fig. 2a). The binding site for Lpm is close to, but does not overlap, the binding site of Myx, Cor, Rip and SQ, which bind to the RNAP switch region, contacting SW1 and the face of SW2 opposite the face contacted by Lpm. The binding site for Lpm is far from the binding sites of other RNAP inhibitors.

**Figure 2.**
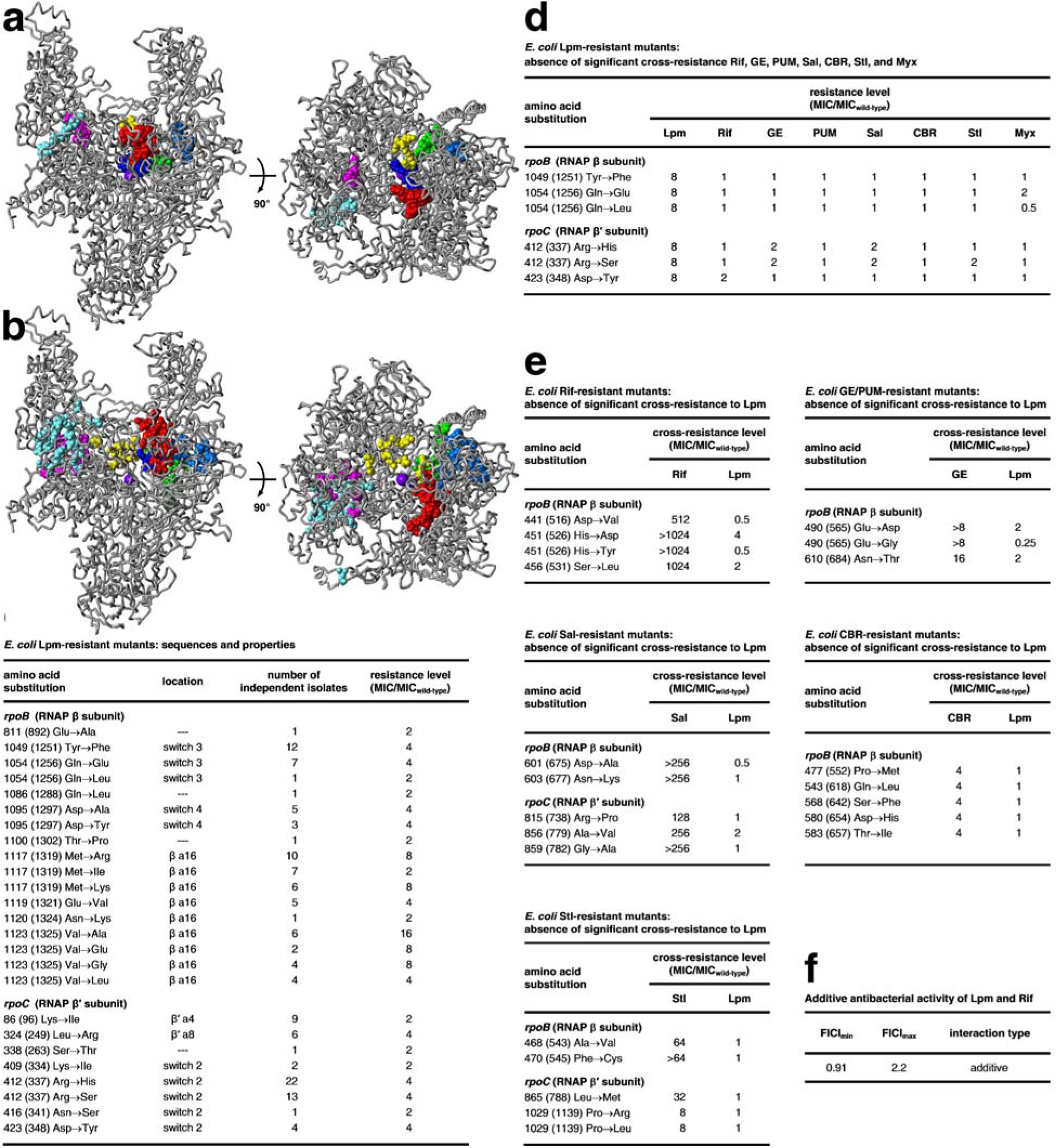
Relationship between binding site and resistance determinant of Lpm and binding sites and resistant determinants of other RNAP inhibitors. **a**, Binding positions of Lpm (cyan; Fig. 1), Rif and Sor (red; PDB 1I6V, 1YNJ, 2A68, 2A69, 4KN4, 4KN7, 4OIR, and 5UHB), GE and PUM (dark blue; PDB 4OIN, 4OIR, and 5X21), CBR and AAP (light blue; PDB 4XSY, 4XSZ, 4ZH2, 4ZH3, 4ZH4, 5UHE, and 5UHF), Sal (green; PDB 4MEX), Stl (yellow; PDB 1ZYR), and Myx and SQ (magenta; PDB 3DXJ, 3EQL, 4YFK, 4YFN, and 4YFX), mapped onto structure of *Mtb* RNAP (gray; two orthogonal views; β’MtbSI and σ omitted for clarity). Violet sphere,2+ 22-23 RNAP active-center Mg. **b**, Resistance determinants of Lpm (cyan; Fig. 2c), Rif and Sor “ (red), GE and PUM^24–25^ (dark blue), CBR703 and AAP^15, 26–27^ (light blue), Sal^28^ (green), Stl^29–30^ (yellow), and Myx, Cor, Rip, and SQ^31–33^ (magenta) mapped onto structure of *Mtb* RNAP. **c**, Sequences and properties of *E. coli* Lpm-resistant mutants residues numbered as in *Mtb* RNAP and, in parentheses, as in *Escherichia coli* RNAP). **d**, Absence of significant cross-resistance of *E. coli* Lpm-resistant mutants to Rif, GE, PUM, Sal, CBR, Stl, and Myx. **e**, Absence of significant cross-resistance of *E. coli* Rif-, GE/PUM-, Sal-, CBR-, and Stl-resistant mutants to Lpm. f, Additive antibacterial activity of Lpm and Rif.

Amino acid substitutions conferring resistance to Lpm were identified by sequencing spontaneous *Bacillus subtilis* Lpm-resistant mutants^7, 9^; additional substitutions were identified by sequencing induced Lpm-resistant mutants of *E. coli* RNAP^9^; and, in this work, a full set of substitutions was identified by extending the latter method (Fig. 2b-c). Lpm-resistant substitutions are obtained at ten sites in RNAP β subunit and seven sites in RNAP β’ subunit (Fig 2b-c). All sites of Lpm-resistant substitutions are located in the immediate vicinity of the Lpm binding site (Fig. 2b), and all high-level (≥4-fold) Lpm-resistant substitutions involve RNAP residues that make direct contact with Lpm or that contact other RNAP residues that, in turn, make direct contact with Lpm (Extended Data Fig. 5a). The resistance determinant for Lpm does not overlap the resistance determinants of other RNAP inhibitors (Fig. 2b). Consistent with the absence of overlap of binding sites and resistance determinants, mutants resistant to Lpm are not cross-resistant to other RNAP inhibitors (Fig. 2d; Extended Data Fig. 6b), and, reciprocally, mutants resistant to other RNAP inhibitors are not cross-resistant to Lpm (Fig. 2e; Extended Data Fig. 6c). Further consistent with the absence of overlap of binding sites and resistance determinants, Lpm and Rif exhibit additive antibacterial activity when co-administered (Fig. 2f).

The RNAP clamp adopts different conformations--ranging from open, to partly closed, to closed--in different crystal structures^14, 16–17^ (Fig. 3a). Differences in clamp conformation arise from rigid-body rotations about a hinge formed by the RNAP switch region^14, 16–17^. Importantly, differences in clamp conformation are observed even in crystal structures of RNAP from a single bacterial species (Fig. 3a; Extended Data Table 2). In the cryo-EM structure of *Mtb* RNAP-Lpm, the clamp adopts an open conformational state, superimposable on the open state in a crystal structure of *Thermus thermophilus (Tth)* RNAP (Extended Data Table 2), but differing by 9° from the partly closed state in a crystal structure of *Tth* RNAP (PDB 5TMC), and differing by 17° from the closed state in crystal structures of *Tth* RPo and *Mtb* RPo (PDB 4G7H and 5UH5) (Fig. 3a-b; Extended Data Fig. 7a). The interactions that Lpm makes with βa16α1, β’a4α1, SW2, SW3, and SW4 require an open clamp conformation (Extended Data Fig. 7b). Each of these regions is positioned to accommodate Lpm in the open state, but not in partly closed and closed states. In particular, βa16α1 and β’a4α1 would severely clash with Lpm in partly closed and closed states (Extended Data Fig. 7). We infer that Lpm, through its interactions with its binding site, induces or stabilizes an open clamp conformation.

**Figure 3.**
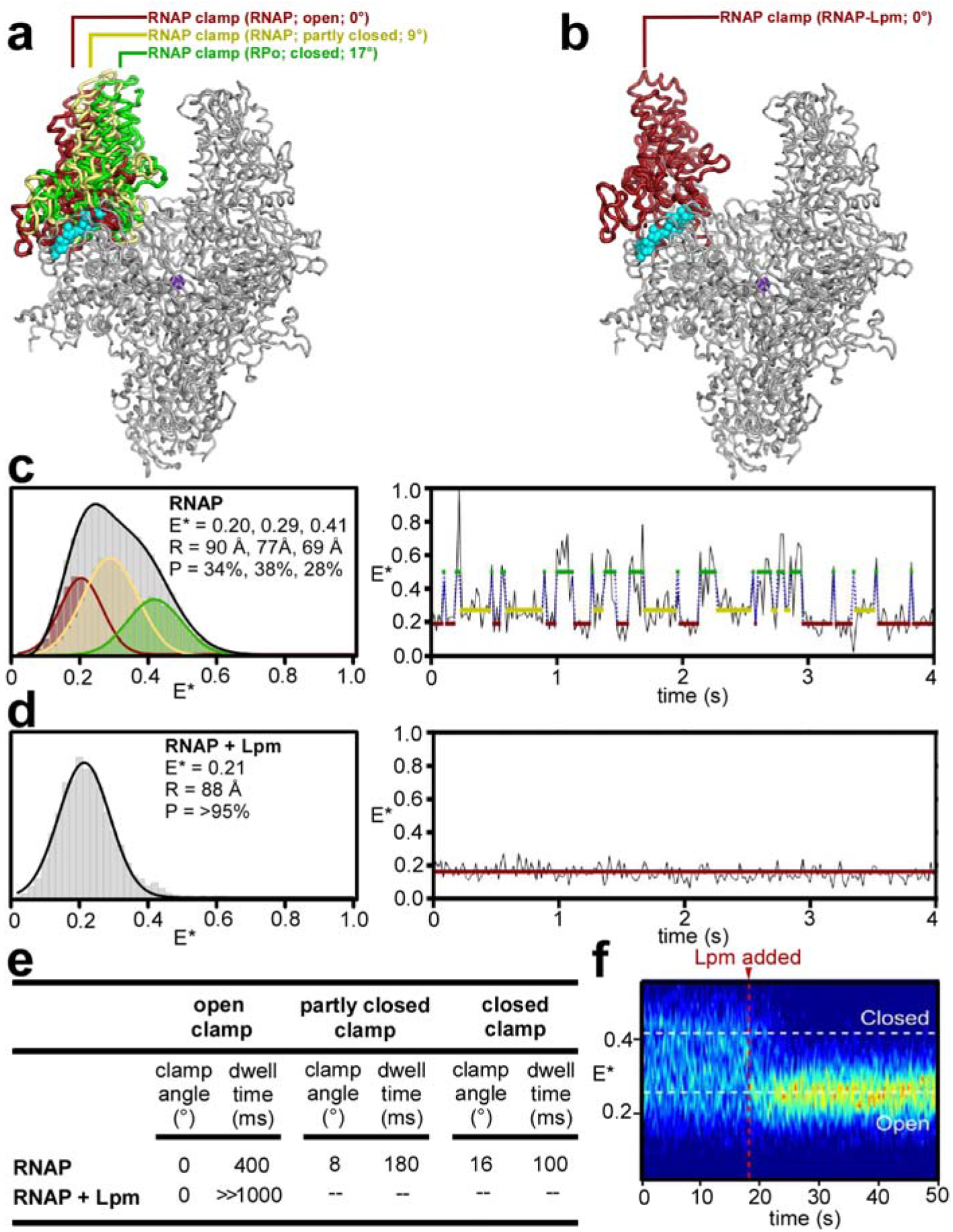
Effects of Lpm on RNAP clamp conformation. **a**, RNAP open (red), partly closed (yellow), and closed (green) clamp conformational states. *Mtb* RNAP-Lpm main mass (view as in Figs. 1–2) and, superimposed, RNAP clamps from crystal structures of *T. thermophilus* RNAP in different crystal lattices (red, yellow, and green; Extended Data Table 2, PDB 5TMC, and PDB 4G7H). **b**, RNAP open-clamp state in *Mtb* RNAP-Lpm. Red, clamp from *Mtb* RNAP-Lpm. Other colors as in a. **c**, Single-molecule FRET data for RNAP holoenzyme in absence of Lpm. Left, histogram. Gray, all states; colors, Hidden Markov Model (HMM)-assigned open, partly closed, and closed states (red, yellow, and green). Right, time trace with HMM-assigned open, partly closed, and closed states (red, yellow, and green). **d**, Single-molecule FRET data for RNAP holoenzyme in presence of Lpm (histogram and time trace as in c). **e**, Summary of RNAP clamp angles and dwell times for open, partly closed, and closed clamp states in absence and presence of Lpm. f, Time trace for RNAP holoenzyme before and after addition of Lpm.

RNAP clamp conformation in solution can be monitored by single-molecule fluorescence resonance energy transfer (FRET) experiments assessing distances between fluorescent probes incorporated specifically at the tips of the two walls of the RNAP active-center cleft^14^ (Fig. 3; Extended Data Fig. 8). Single-molecule FRET experiments analyzing RNAP holoenzyme in the absence of Lpm show a broad multimodal distribution of FRET efficiencies, indicative of open, partly closed, and closed states (Fig. 3c-left, gray curve). In contrast, analogous experiments in the presence of Lpm show a narrow unimodal peak with a FRET efficiency indicative of the open state (Fig. 3d, left). We conclude that Lpm traps the open-clamp state in solution. Single-molecule time traces in the absence of Lpm show static and dynamic molecules; the dynamic molecules (34%; N = 207) show transitions between three distinct FRET efficiency levels--corresponding to open-clamp, partly-closed-clamp, and closed-clamp states, with dwell times on the 100-400 ms time scale--indicating dynamic transitions between interconverting clamp states (Fig. 3c-left, colored curves, 3c-right, and 3e; Extended Data Fig. 8b-d). In contrast, analogous single-molecule time traces in the presence of Lpm show only the FRET efficiency level corresponding to the open-clamp state, indicating stable trapping of the open-clamp state (Fig. 3d, right, and 3e; Extended Data Fig. 8b-d). Single-molecule time traces before and after addition of Lpm show a rapid (<2 s) transition from the multimodal, dynamic clamp profile of RNAP in the absence of Lpm to the unimodal, non-dynamic clamp profile of RNAP in the presence of Lpm, demonstrating rapid trapping of the open-clamp state (Fig. 3f). We conclude, based both on cryo-EM structure and on FRET, that Lpm traps the open-clamp state.

Strikingly, the effect of Lpm on RNAP clamp conformation is *opposite* the effects of Myx, Cor, and Rip on RNAP clamp conformation. Lpm, through interactions with RNAP switch-region SW2, SW3, and SW4, traps an open-clamp state^14^; in contrast, Myx, Cor and Rip, through interactions with SW1 and the opposite face of SW2, trap a closed-clamp state (Fig. 3). The different effects of these inhibitors on RNAP clamp conformation presumably underlie their different effects on isomerization of RPc to RPo (inhibition of an early step in isomerization by Lpm; inhibition of a later step in isomerization by Myx, Cor, and Rip^10–11^).

The binding site on RNAP for Lpm overlaps the proposed binding site on RNAP for pause-inducing and termination-inducing RNA hairpins^34^. It is attractive to speculate that the Lpm binding site is an “allosteric trigger” for RNAP clamp opening and that ligands of this site--including not only Lpm but also pause-inducing and termination-inducing RNA hairpins--trigger RNAP clamp opening by binding to the site.

The effects of Lpm on RNAP clamp conformation suggest a model for the mechanism by which Lpm inhibits isomerization of RPc to RPo^8–11^ in transcription initiation (Fig. 4). Because σR2 (the σ module that recognizes promoter -10 elements^35–39^) interacts with the RNAP clamp and σR4 (the σ module that recognizes promoter -35 elements^35, 37–39^ “) interacts with the main mass of RNAP, differences in RNAP clamp conformation necessarily result in differences in the spatial relationship of σR2 and σR4 (Fig. 4, left panels). We propose that, in the closed-clamp state, σR2 and σR4 can engage simultaneously with promoter -10 and -35 elements (Fig. 4a, middle and right panels), but, in the open-clamp state, σR2 and σR4 cannot engage simultaneously with promoter -10 and -35 elements (Fig. 4b, middle and right panels). In particular, we propose that the σR2 “Trp wedge,” which intercalates into DNA and stacks on the non-template-strand nucleotide at promoter position -12, nucleating promoter unwinding in formation of RPo^38–39^, can engage DNA only in the closed-clamp state, and that the σR2 “NT-11 pocket,” which captures an unstacked non-template-strand nucleotide at promoter position -11, stabilizing promoter unwinding upon formation of RPo^36–39^, can engage DNA only in the closed-clamp state (Fig. 4a, middle and right panels). The proposal that Lpm inhibits transcription initiation by trapping an open-clamp state, thereby preventing simultaneous engagement of -10 and -35 elements, is consistent with recently proposed models for transcription initiation.^40^

**Figure 4.**
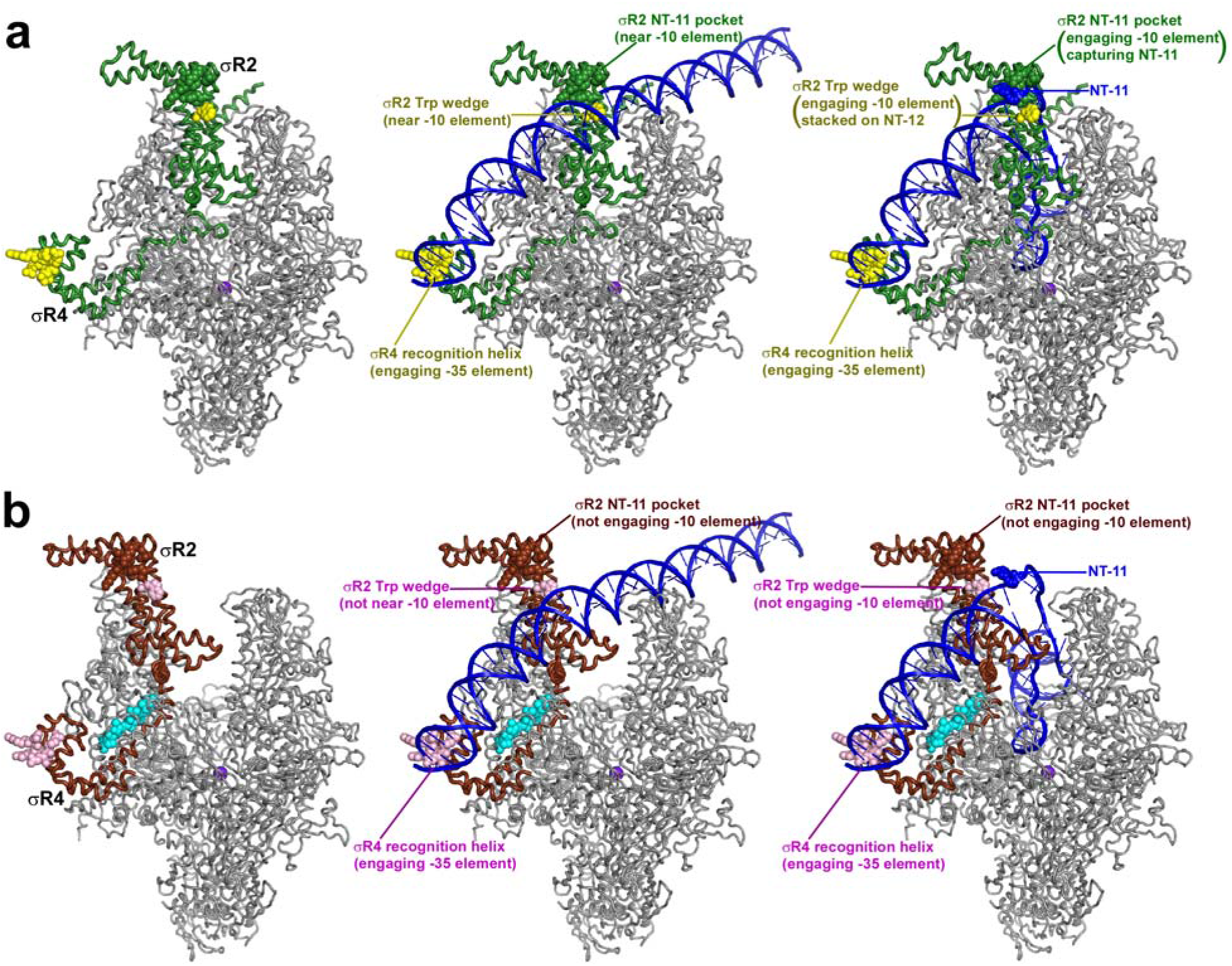
Proposed mechanism of transcription inhibition by Lpm. **a**, RNAP holoenzyme (left), RPc (center; DNA as in^35^), and RPo (right, DNA as in^37–39^) for RNAP in absence of Lpm (closed clamp). σR2 and σR4 simultaneously engage promoter -10 and -35 elements in RPc and RPo. Atomic coordinates from PDB 5UH5. Green ribbons, σ; green surface, σR2 NT-11 pocket; yellow surfaces, σR2 Trp wedge and σR4 recognition helix; blue, DNA. Other colors as in Figs. 1–2. **b**, As a, but for RNAP in presence of Lpm (open clamp). σR2 and σR4 cannot simultaneously engage promoter -10 and -35 elements in RPc and RPo. Brown ribbons, σ; brown surface, σR2 NT-11 pocket; pink surface, σR2 Trp wedge and σR4 recognition helix; cyan, Lpm. Other colors as in **a**.

The methods and results of this report provide a strategy to obtain structural information for *any* RNAP inhibitor, irrespective of species selectivity and effects on RNAP conformation. The use of cryo-EM to determine the structure of RNAP-Lpm overcame obstacles that had precluded structure determination by x-ray crystallography: (1) the species selectivity of Lpm (which does not inhibit *Thermus* RNAP) and (2) the trapping by Lpm of an RNAP open-clamp conformation (which is incompatible with known crystal lattices of *non-Thermus* bacterial RNAP).

The results of this report provide fundamental information on the binding site and mechanism of Lpm and provide insights for antibacterial drug development involving Lpm. The primary limitation of Lpm as an antibacterial drug is that Lpm has no systemic bioavailability and thus is ineffective against systemic infections, such as tuberculosis and *S. aureus* blood and lung infections^1, 2, 5^. The chemical strategy and synthetic procedure demonstrated here for appending substituents at the Lpm C4” hydroxyl (Fig S5; a1) provides a means to overcome this limitation. It should be possible to append substituents improving physical and pharmacokinetic properties, nanoparticle drug-delivery modules, or even other antibacterial-agent pharmacophores at the Lpm C4” hydroxyl while retaining RNAP-inhibitory activity.

## Acknowledgements

This work was supported by NIH grants GM041376, AI104660, and AI109713-01 to R.H.E. We thank W. Fenical, H. Irschik, R. Jansen, A. L. Sonenshein, and E. Steinbrecher for compounds; J. Mukhopadhyay, S. Rodrigue, and J. Swezey for strains and plasmids; APS at Argonne National Laboratory for beamline access; the Rutgers University cryo-EM facility and the National Resource for Automated Molecular Microscopy (supported by NIH grant GM103310 and Simons Foundation grant 349247) for cryo-EM access; and K. Callanan, B. Carragher, and C. Potter for discussion.

## Data Availability

Atomic coordinates and structure factors for reported structures have been deposited with the Electron Microscopy Data Bank and Protein Data Bank under accession codes EMDB 4230 and PDB 6FBV *(Mtb* RNAP-Lpm) and PDB 6ASG *(T. thermophilus* RNAP core enzyme).

## Author Contributions

W.L. and A.M prepared RNAP derivatives. W.L., K.D., Y.L, R.Y., Z.Z., E.E., and D.T. performed structure determination. D. Degen, R.Y.E., E.S., M.G., A.S., S.M., and Y.J. assessed RNAP-inhibitory activities, antibacterial activities, resistance, and cross-resistance. A.M., D. Duchi, and A.K. assessed FRET. Y.W.E., S.D., H.Z., and C.Z. prepared RNAP inhibitors. R.H.E. designed the study, analyzed data, and wrote the paper.

## Author Information

R.H.E. has patent filings. The other authors declare that no competing interests exist. Correspondence and requests for materials should be addressed to R.H.E.

## METHODS

### Lpm

Lpm was the kind gift of A.L. Sonenschein (Tufts University, Medford MA), was prepared as described^41^, or was purchased from BioAustralis or Biorbyt.

### Lpm analogs: previously described Lpm analogs

Lpm analog “a4” (3”-bromo-3”,5”-dideschloro-Lpm; compound 3 of ref. 19) was prepared as described^19^. Lpm analogs “a5” (69-1d), “a6” (70-9d), “a7” (70-10d), “a8” (70-11d), “b2” (53-11d), “c1” (54-4d), and “e1” (54-3d) were prepared as described^20–21^. Lpm analog “b1” (OP-1118^18^) was purchased from Toronto Research Chemicals.

### Lpm analogs: Lpm analogs “a2” (5”-deschloro-Lpm) and “d1” (desaryl-Lpm)

Lpm analogs “a2” (5”-deschloro-Lpm) and “d1” (desaryl-Lpm) were prepared by fermentation of the Lpm producer strain *Dactylosporangium aurantiacum* hamdenensis NRRL 18085^42^ (ARS Patent Culture Collection, Peoria IL) in growth media containing limiting chloride [growth media with KCl replaced by KBr; residual chloride concentration = 40 μM, quantified using QuantiChromT Chloride Assay Kit (BioAssays Systems)]. First-stage (0.2 L) and second-stage (10 L) cultures were prepared as described^19^, and second-stage culture broths were harvested as described^19^. Culture broths were adjusted to pH 7 by addition of 1 ml 12 N NaOH, supplemented with 5 L acetone, shaken 1 h at 22°C, and extracted with 3x4 L ethyl acetate. The extracts were evaporated, and the resulting material (20 g) was re-dissolved in 300 ml ethyl acetate, filtered through Whatman #1 filter paper (Sigma-Aldrich), and evaporated. The resulting material (14 g) was partitioned in 900 ml 1:1:1 (v/v/v) chloroform-methanol-water, and the lower phase was evaporated. The resulting material (11 g) was chromatographed on silica gel (Sigma-Aldrich; 3 cm x 30 cm; stepwise elution with 2-50% methanol in chloroform, followed by elution with 100% methanol). UV-absorbant (254 nm) fractions were collected, evaporated, and assayed for antibacterial activity by spotting on H-top-agar/LB-agar^43^ plates seeded with 10^9^ cfu *E. coli* D21f2tolC^44^, incubating 16 h at 37, and assessing growth inhibition. Active fractions eluting at ~5% to ~10% methanol were pooled, further purified by re-chromatography on silica gel (procedures essentially as above), further purified by reversed-phase HPLC [Hitachi 7450 with L7450 detector; Phenomenex C18 Jupiter semi-prep; 1:1 (v/v) acetonitrile-water isocratic elution; flow rate = 2 ml/min], and evaporated, yielding 80 mg Lpm analog “a3” [MALDI-MS m/z: calculated 1011.41 (M + Na+); found 1011.07] and 8 mg Lpm analog “a2” [MALDI-MS m/z: calculated 1045.59 (M + Na+); found 1044.60, 1046.70]. Material eluting at 100% methanol was further purified by reversed-phase HPLC (procedures essentially as above), and evaporated, yielding 0.2 mg Lpm analog “d1” [MALDI-MS m/z: calculated, 846.98 (M + Na^+^); found, 847.90].

### Lpm analogs: Lpm analog “a1” (4”-O-benzyl-Lpm)

Lpm analog “a1” (4”-O-benzyl-Lpm) was prepared from Lpm by semi-synthesis as follows: To Lpm (Biorbyt: 3 mg; 2.8 μmol) in 50 μl anhydrous dimethylformamide (Sigma-Aldrich), was added anhydrous potassium carbonate (Thermo-Fisher: 3 mg, 22 μmol), and the reaction mixture was stirred 30 min at 50°C and then allowed to cool to room temperature. An aliquot (10 μl) of 0.28 M benzyl bromide in anhydrous dimethylformamide [2.8 μmol; prepared by dissolving 3.4 μl benzyl bromide (Sigma-Aldrich) in 50 μl anhydrous dimethylformamide immediately before use] was added, and the reaction mixture was stirred 30 min at 50°C. The reaction mixture was evaporated to remove dimethylformamide and then re-suspended in 600 μl 100 mM monobasic sodium phosphate (Thermo-Fisher). Precipitated solids were collected by centrifugation, rinsed with water, re-dissolved in methanol, and purified by reversed-phase HPLC (Phenomenex C18 Luna, 100 Å, 25 mm x 4.6 mm; phase A = water; phase B = acetonitrile; gradient = 40% B at 0 min, 100% B at 20 min; flow rate = 1 ml/min). Lpm analog “a1” eluted at 20 minutes. Yield: 0.35 mg, 10%. MALDI-MS m/z: calculated, 1171.44 (M + Na+); found 1169.64, 1171.64.

### Other RNAP inhibitors

Cor A and Rip A were the kind gift of H. Irschik and R. Jansen (Helmholtz Institut, Braunschweig, Germany), Stl was the kind gift of E. Steinbrecher (Upjohn-Pharmacia, Kalamazoo, MI), and Sal A was the kind gift of W. Fenical (The Scripps Research Institute, La Jolla, CA). Myx B was prepared as described^45^, SQ1 (compound 1 of ref. 46) was prepared as described^46^, GE was prepared as described^47^, and PUM was prepared as described^25^. Rif was purchased from Sigma-Aldrich, and CBR was purchased from Maybridge.

### *M. tuberculosis* RNAP σ^A^ holoenzyme

For all experiments other than the experiments in Extended Data Fig. 3d, *Mtb* RNAP σ^A^ holoenzyme was prepared as described^15^.

For experiments in in Extended Data Fig. 3d, *Mtb* RNAP σ^A^ holoenzyme and *Mtb* RNAP σ^A^ holoenzyme derivatives were prepared by the same procedure, but using plasmid pCOLADuet-Mtb-rpoBC [prepared by replacing the NcoI-BamHI segment of plasmid pCOLA-Duet (Novagen) by the NcoI-BamHI segment of a DNA fragment carrying CCATGGTG followed by *Mtb rpoB* codons 3-1174 followed by TAAGGATCC, prepared by PCR using as template plasmid JF09^48^ (gift of S. Rodrigue, Universitė de Sherbrooke, Canada), and then replacing the NdeI-MfeI segment of the resulting plasmid by the NdeI-MfeI segment of a DNA fragment carrying CATATG followed by *Mtb rpoC* codons 2-1316 followed by TAGCAATTG, prepared by PCR using as template plasmid JF10^48^ (gift of S. Rodrigue, Universitė de Sherbrooke, Canada)], or derivatives thereof constructed using site-directed mutagenesis (QuikChange Site-Directed Mutagenesis Kit; Agilent), in place of plasmid pCOLA-rpoB-rpoC^49^.

### *S. aureus* RNAP σ^A^ holoenzyme

*S. aureus* RNAP σ^A^ holoenzyme was prepared as described^25^.

### *E. coli* RNAP σ^70^ holoenzyme

For experiments in Extended Data Fig. 5, hexahistidine-tagged *E. coli* RNAP σ^70^ holoenzyme was prepared from *E. coli* strain XE5 4^50^ transformed with plasmid pREII-NHα^50^, using procedures as described^51^

For experiments in Figs. 3c-e and Extended Data Fig. 8c-d, fluorescent-probe-labelled, hexahistidine-tagged *E. coli* RNAP σ^70^ holoenzyme was prepared using unnatural-amino-acid mutagenesis^14, 52^ of co-expressed genes encoding RNAP β’, β, α, and ω subunits and σ^70^, yielding an RNAP σ^70^ holoenzyme derivative containing 4-azido-L-phenylalanine (AzF) at position 284 of β’ and position 106 of β, followed by azide-specific Staudinger ligation^14, 53–55^ to incorporate the fluorescent probes Cy3B and Alexa647 at position 284 of β’ and position 106 of β, as follows (Extended Data Fig. 8a): Single colonies of *E. coli* strain BL21(DE3) (Millipore) co-transformed with plasmid pVS10-rpoB106am;rpoC284am [constructed by use of site-directed mutagenesis (QuikChange Site-Directed Mutagenesis Kit; Agilent) to replace *rpoB* codon 106 and *rpoC* codon 284 by amber codons in plasmid pVS10^56^], plasmid pRSFsigma70^57^, and plasmid pEVOL-pAzF^52^ were used to inoculate 20 ml LB broth^43^ containing 100 μg/ml ampicillin, 50 μg/ml kanamycin, and 35 μg/ml chloramphenicol, and cultures were incubated 16 h at 37°C with shaking. Culture aliquots (2x10 ml) were used to inoculate LB broth^43^ (2x1 L) containing 2 mM AzF (Chem-Impex International), 100 μg/ml ampicillin, 50 μg/ml kanamycin, and 35 μg/ml chloramphenicol, cultures were incubated with shaking until OD_600_ = 0.6; L-arabinose was added to 0.2% and IPTG was added to 1 mM, and cultures were further incubated 16 h at 16°C with shaking. Cells were harvested by centrifugation (4,000 x g; 20 min at 4°C), re-suspended in 20 ml buffer A (10 mM Tris-HCl, pH 7.9, 200 mM NaCl, and 5% glycerol), and lysed using an EmulsiFlex-C5 cell disrupter (Avestin). The lysate was cleared by centrifugation (20,000 x g; 30 min at 4°C), precipitated with polymin P (Sigma-Aldrich) as described^51^, and precipitated with ammonium sulfate as described^51^. The sample was dissolved in 30 ml buffer A and loaded onto a 5 ml column of Ni-NTA-agarose (Qiagen) pre-equilibrated in buffer A, and the column was washed with 50 ml buffer A containing 10 mM imidazole and eluted with 25 ml buffer A containing 200 mM imidazole. The sample was further purified by anion-exchange chromatography on Mono Q 10/100 GL (GE Healthcare; 160 ml linear gradient of 300-500 mM NaCl in 10 mM Tris-HCl, pH 7.9, 0.1 mM EDTA, and 5% glycerol; flow rate = 2 ml/min). Fractions containing AzF-derivatized hexahistidine-tagged *E. coli* RNAP σ^70^ holoenzyme were pooled, concentrated to ~1 mg/ml using 30 kDa MWCO Amicon Ultra-15 centrifugal ultrafilters (Millipore), and stored in aliquots at -80°C. A reaction mixture containing 10 μM AzF-derivatized hexahistidine-tagged *E. coli* RNAP σ^70^ holoenzyme, 100 μM Alexa647-pentanoyl-ethylenediaminyl-phosphine (prepared as described^54–55^), and 100 μM Cy3B-carboyl-ethylenediaminyl-phosphine (prepared as described^54–55^) in 1 ml buffer B (50 mM Tris-HCl, pH 7.9, 100 mM KCl, 5% glycerol, and 2% dimethylformamide) was incubated 1 h at 15 °C, incubated 16 h on ice, and subjected to 5 cycles of buffer exchange (dilution with 5 ml buffer B, followed by concentration to 0.5 ml) using 30 kDa MWCO Amicon Ultra-15 centrifugal ultrafilters (Millipore). The sample was further purified by gel-filtration chromatography on HiLoad 16/60 Superdex 200 prep grade (GE Healthcare) pre-equilibrated in buffer C (20 mM Tris-HCl, pH 8.0, 100 mM NaCl, 5 mM MgCl2, 1 mM β-mercaptoethanol, and 5% glycerol) and eluted in buffer C. Fractions containing fluorescent-probe-labelled hexahistidine-tagged *E. coli* RNAP σ^70^ holoenzyme were pooled, concentrated to 1 mg/ml in buffer C using 30 kDa MWCO Amicon Ultra-15 centrifugal ultrafilters (Millipore) and stored in aliquots at -80°C.

For experiments in Fig. 3f, fluorescent-probe-labelled, hexahistidine-tagged, FLAG-tagged *E. coli* RNAP σ^70^ holoenzyme was prepared using unnatural-amino-acid mutagenesis^52^ of genes encoding RNAP β’ and β subunits to incorporate azidophenylalanine at position 284 of β’ and position 106 of β, azide-specific Staudinger ligation^14, 53–55^ to incorporate Cy3B at position 284 of the resulting β’ derivative and Alexa647 at position 106 of the resulting β derivative, and *in vitro* reconstitution of RNAP^14,55,58^ from the resulting β’ and β derivatives, RNAP α and ω subunits, and σ^70^, as described^14, 54–55^.

Efficiencies of incorporation of fluorescent probes were determined from UV/Vis-absorbance measurements and were calculated as:
concentration of product = [A_280_ - ε_Cy3B,280_(A_Cy3B,559_/ε_Cy3B,559_) - ε_Alexa,280_(A_Alexa,652_/ε_Alexa647,652_)]/ε_P,280_

Cy3B labelling efficiency = 100%[(A_Cy3B,559_/ε_Cy3B 559_)/(concentration of product)]

Alexa647 labelling efficiency = 100%[(A_Alexa,652_/ε_Alexa,652_)/(concentration of product)]

where A_280_ is the measured absorbance at 280 nm, A_Cy3B,559_ is the measured absorbance at the long-wavelength absorbance maximum of Cy3B (559 nm), A_Alexa,652_ is the measured absorbance at the long-wavelength absorbance maximum of Alexa647 (652 nm), ε_P,280_ is the molar extinction coefficient of RNAP σ^70^ holoenzyme at 280 nm (240,000 M^−1^ cm^−1^), ε_Cy3B,280_ is the molar extinction coefficient of Cy3B at 280 nm (7,350 M^−1^ cm^−1^), B_Alexa,280_ is the molar extinction coefficient of Alexa647 at 280 nm (10,400 M^−1^ cm^−1^), and ε_Cy3B,559_ is the extinction coefficient of Cy3B at its long-wavelength absorbance maximum (130,000 M^−1^ cm^−1^), and B_Alexa,652_ is the extinction coefficient of Alexa647 at its long-wavelength absorbance maximum (245,000 M^−1^ cm^−1^). Labelling efficiencies were ~90% for Cy3B and ~70% for Alexa647.

Specificities of incorporation of fluorescent probes were determined from the observed labelling efficiencies of (i) the labelling reaction with the AzF-derivatized hexahistidine-tagged *E. coli* RNAP σ^70^ holoenzyme and (ii) a control labelling reaction with non-AzF-derivatized hexahistidine-tagged *E. coli* RNAP σ^70^ holoenzyme, and were calculated as:
labelling specificity = 100%[1 - [(labelling efficiency with P)/(labelling efficiency with AzF-P)]

where AzF-P is AzF-derivatized hexahistidine-tagged *E. coli* RNAP σ^70^ holoenzyme, and P is non-AzF-derivatized hexahistidine-tagged *E. coli* RNAP σ^70^ holoenzyme. Labelling specificities were >90%.

### *T. thermophilus* RNAP core enzyme

*T. thermophilus* RNAP core enzyme was prepared as described^36^.

### RNAP-inhibitory activities

Fluorescence-detected RNAP-inhibition assays with the profluorescent substrate γ-[2’-(2-benzothiazoyl)-60-hydroxybenzothiazole]-ATP (BBT-ATP; Jena Bioscience) were perfomed as described^32^, using 75 nM RNA holoenzyme (*Mtb* RNAP σ^A^ holoenzyme or derivative, *S. aureus* σ^A^ RNAP holoenzyme, or *E. coli* RNAP σ^70^ holoenzyme) and 20 nM DNA fragment containing bacteriophage T5 N25 promoter (prepared as described^24^).

Radichemical RNAP-inhibition assays with HeLa nuclear extract (human RNAP I/II/III) were performed as described^28.^

Half-maximal inhibitory concentrations (IC50s) were calculated by non-linear regression in SigmaPlot (Systat Software).

### Lpm-resistant mutants

Lpm-resistant mutants were isolated using procedures analogous to those used for isolation of Myx-resistant mutants in ref. 31. Random mutagenesis of *E. coli rpoC* plasmid pRL663^59^ and *E. coli rpoB* plasmid pRL706^60^ was performed by use of PCR amplification, exploiting the baseline error rate of PCR amplification. Mutagenesis reactions were performed using the QuikChange Site-Directed Mutagenesis Kit (Agilent) with plasmid pRL663 and oligodeoxyribonucleotide forward and reverse primers corresponding to nucleotides 1-20, 67-68, 77-81, 93-100, 245-256, 259-265, 325-355, 378-382, 393-403, 425-433, 466-481, 1319-1327, and 1347-1360 of *E. coli rpoC*, or with plasmid pRL706 and oligodeoxyribonucleotide forward and reverse primers corresponding to nucleotides 854-857, 890-899, 914-922, and 1246-1342 of *E. coli rpoB* (primers at 75 nM; all other components at concentrations as specified by the manufacturer). Mutagenized plasmid DNA was introduced by transformation into *E. coli* XL1-Blue (Agilent). Transformants (~10^4^ cells) were applied to LB-agar^43^ plates containing 200 μg/ml ampicillin, plates were incubated 16 h at 37°C, and plasmid DNA was prepared from the pooled resulting colonies. The resulting pooled mutagenized plasmid DNA was introduced by transformation into uptake-proficient, efflux-deficient *E. coli* strain D21f2tolC^44^. Transformants (~10^3^ cells) were applied to LB-agar^43^ plates containing 5 μg/ml Lpm (the minimal concentration that prevents colony formation by wild-type transformants), 200 μg/ml ampicillin, and 1 mM IPTG, and plates were incubated 24-48 h at 37°C. Lpm-resistant mutants were identified by the ability to form colonies on this medium, were confirmed by re-streaking on the same medium, and were demonstrated to contain plasmid-linked Lpm-resistant mutations by preparing plasmid DNA, transforming *E. coli* D21f2tolC with plasmid DNA, and plating transformants on the same medium. Nucleotide sequences of *rpoB* and *rpoC* were determined by Sanger sequencing (nine primers per gene).

### Resistance and cross-resistance levels

Experiments in Fig. 2c and Extended Data Fig. 6b-c assessing resistance and cross-resistance levels of Lpm-resistant *E. coli* D21f2tolC pRL706 and *E. coli* D21f2tolC pRL663 derivatives (preceding section) and Myx/Cor/Rip-resistant *E. coli* D21f2tolC pRL706 and *E. coli* D21f2tolC pRL663 derivatives^31^ were performed using spiral gradient endpoint assays^61–63^ on 150 mm x 4 mm exponential-gradient plates containing LB agar^43^, 0.05-50 μg/ml test compound (Lpm, Rif, Myx, Cor, or Rip), 200 mg/ml ampicillin, and 1 mM IPTG. Test compounds were applied to plates using an Autoplate 4000 spiral plater (Spiral Biotech). Single colonies of transformants of *E. coli* D21f2tolC^44^ were inoculated into 5 ml LB broth^43^ containing 200 μg/ml ampicillin and incubated at 37°C with shaking until OD_600_ = 0.4-0.6, IPTG was added to 1 mM, and cultures were incubated 1 h at 37°C with shaking.

Diluted aliquots (~1x10^9^ cfu/ml for Fig. 2c and ~1x10^8^ cfu/ml for Extended Data Fig. 6b) were swabbed radially onto plates, and plates were incubated 16 h at 37°C. For each culture, the streak length was measured using a clear plastic template (Spiral Biotech), the test-compound concentration at the streak endpoint was calculated using the program SGE (Spiral Biotech), and the minimum inhibitory concentration (MIC) was defined as the test-compound concentration at the streak endpoint.

Experiments in Fig. 2d assessing resistance and cross-resistance levels of chromosomal Lpm-resistant mutants [mutations transferred from pRL706 or pRL663 derivatives of preceding section to chromosome of *E. coli* D21f2tolC^44^ by λ-Red-mediated recombineering (procedures essentially as described^64^, but using transformation rather than electroporation)] were performed using broth microdilution assays^65^. Single colonies were inoculated into 5 ml LB broth^43^ and incubated at 37°C with shaking until OD_600_ = 0.4-0.8. Diluted aliquots (~5x10^4^ cells) in 97 μl LB broth were dispensed into wells of a 96-well plate, and were supplemented with 3 μl methanol or 3 μl of a 2-fold dilution series of Lpm (MIC_wild-type_ = 1.56 μg/ml), Rif (MIC_wild-type_ = 0.20 μg/ml), CBR (MIC_wild-type_ = 6.25 μg/ml), Sal (MIC_wild-type_ = 0.049 μg/ml), Stl (MIC_wild-type_ = 3.13 μg/ml), or Myx (MIC_wild-type_ = 0.20 μg/ml), in methanol (final concentrations = 0 and 0.006-50 μg/ml), or diluted aliquots (~1x10^5^ cells) in 50 μl LB broth were supplemented with 50 μl LB broth or 50 μl of a 2-fold dilution series of GE (MIC_wild-type_ = 500 μg/ml) or PUM (MIC_wild-type_ = 400 μg/ml) in LB broth (final concentrations = 0 and 25-2000 μg/ml). Plates were incubated 16 h at 37°C. The MIC was defined as the lowest tested concentration that inhibited bacterial growth by ≥90%.

Experiments in Fig. 2e assessing Lpm-cross-resistance levels of Rif-, GE/PUM-, and Sal-resistant mutants were performed as described for experiments in Fig. 2d, but analyzing a panel of *E. coli* D21f2tolC derivatives^25, 28^ comprising 4 chromosomal Rif-resistant mutants, 3 chromosomal GE/PUM-resistant mutants, and 5 chromosomal Sal-resistant mutants.

Experiments in Fig. 2e assessing Lpm-cross-resistance levels of CBR- and Stl-resistant mutants were performed essentially as described for experiments in Fig. 2d, but analyzing a panel of *E. coli* D21f2tolC pRL706 and *E. coli* D21f2tolC pRL663 derivatives^26, 29^ comprising 5 CBR-resistant mutants and 5 Stl-resistant mutants. Single colonies were inoculated into 5 ml LB broth^43^ containing 200 μg/ml ampicillin and incubated at 37°C with shaking until OD_600_ = 0.4-0.6, IPTG was added to 1 mM, and cultures were incubated 1 h at 37°C with shaking. Diluted aliquots (~5x10^4^ cells) in 97 μl LB broth containing 200 μg/ml ampicillin and 1 mM IPTG were dispensed into wells of a 96-well plate, and were supplemented with 3 μl methanol or 3 μl of a 2-fold dilution series of CBR or Stl in methanol and further processed as described for experiments in Fig. 2d.

### Checkerboard interaction assays

Antibacterial activities of combinations of Lpm and Rif were assessed in checkerboard interaction assays^70–71^. Broth-macrodilution (procedures essentially as described^65^) were performed in checkerboard format, using *E. coli* D21f2tolC^44^ and using LB broth^43^ containing pairwise combinations of: (i) Lpm at 2.0x, 1.75x, 1.5x, 1.3125x, 1.25x, 1.125x, 0.9375x, 0.75x, 0.5625x, 0.5x, 0.375x, 0.25x, and 0.1875x MIC_Lpm_ and (ii) Rif at 2.0x, 1.75x, 1.5x, 1.3125x, 1.25x, 1.125x, 0.9375x, 0.75x, 0.5625x, 0.5x, 0.375x, 0.25x, and 0.1875x MIC_Rif_. Fractional inhibitory concentrations (FICs), FIC indices (FICIs), and minimum and maximum FICIs (FICI_min_ and FICI_max_) were calculated as described^71^. FICI_min_ < 0.5 was deemed indicative of super-additivity (synergism), FICI_min_ > 0.5 and FICI_max_ ≤ 4.0 was deemed indicative of additivity, and FICI_max_ > 4.0 was deemed indicative of sub-additivity (antagonism)^70–71^.

### Cryo-EM structure determination *(M. tuberculosis* RNAP-Lpm): sample preparation

Lacey carbon grids (LC300-CU-100; Electron Microscopy Sciences) were glow-discharged for 30 s using a glow-discharge cleaning system (PELCO easiGlow; Ted Pella) and mounted in the sample chamber of an EM GP grid plunger (Leica) at 18°C and relative humidity = 95%. Grids were spotted with 3.5 μl 1 μM *Mtb* RNAP-Lpm and 50 μM Lpm in 20 mM Tris-HCl, pH 8.0, 75 mM NaCl, 5 mM MgCl_2_, 5 mM dithiothreitol, and 0.1% n-octyl-β-D-glucopyranoside [prepared by pre-equilibrating 150 μl samples containing components other than n-octyl-β-D-glucopyranoside (Biosynth) 30 min at 25°C, and adding n-octyl-β-D-glucopyranoside immediately before spotting], incubated 10 s, blotted with filter paper (Whatman Grade 541; Sigma-Aldrich) for 2.3 s, flash-frozen by plunging in liquid ethane cooled with liquid nitrogen, and stored in liquid nitrogen.

### Cryo-EM structure determination (*M. tuberculosis* RNAP-Lpm): data collection and data reduction

Data were collected at the National Resource for Automated Molecular Microscopy of the Simons Electron Microscopy Center using a 300 keV Titan Krios (FEI/ThermoFisher) electron microscope equipped with a K2 Summit direct electron detector (Gatan) operating in counting mode and a GIF Quantum energy filter (Gatan) with slit width of 20 eV. Data were collected semi-automatically using the software package Leginon^72^, a nominal magnification of 130,000x, a calibrated pixel size of 1.061 Å, and a dose rate of 8 electrons/pixel/s. Movies were recorded at 200 ms/frame for 10 s (50 frames total), resulting in a total radiation dose of 72.05 electrons/Å^2^ per movie Defocus range was varied between 1.0 μm and 2.0 μm. A total of 2,458 micrographs were recorded from two grids over 2 days.

Data were processed as summarized in Extended Data Fig. 1b-d. Data processing was performed on an Ubuntu 16.04 Linux GPU workstation (Titan Computers) containing four GTX 1080 Ti graphic cards (Nvidia)^73^. Frames in individual movies were aligned using MotionCor2^74^, and the first 35 frames per movie were merged to calculate individual micrographs. Contrast-transfer-function estimations were performed using Gctf^75^, yielding defocus-range estimates ranging from 0.4 μm to 4.0 μm for individual micrographs. Initial particle picking was performed on 25 selected micrographs from the first grid (8,609 particles) using the Xmipp routine of the software package Scipion v1.1^76^, and particles were used for two-dimensional class averaging in Relion v2.0.5^77^. Eight distinct two-dimensional classes were selected as templates for picking 452,912 particles from 1,236 selected micrographs from the first grid and 366,594 particles from 1,025 selected micrographs from the second grid, using the Autopick routine of Relion. Two- and three-dimensional classifications were performed on the 452,912 particles from the first grid and 366,594 particles from the second grid, using Relion and using a 60 Å low-pass-filtered map calculated from the crystal structure of *Mtb* RPo (PDB 5UH5^15^; protein residues only) as the starting reference model for three-dimensional classification. Following identification and removal of heterogeneous particles in the two- and three-dimensional classifications, independent but similar density maps were obtained from 126,977 particles from the first grid and 97,412 particles from the second grid. Following merging of the 224,389 particles from the first and second grids, three-dimensional auto-refinement using Relion, and local angular sampling using Relion, a final density map was obtained from a subset of 68,895 particles. Gold-standard Fourier-shell-correlation analysis (FSC^78^) indicated a mean map resolution of 3.52 Å, and ResMap^79^ indicated a median map resolution of 3.5 Å (Extended Data Fig. 1e-f; Extended Data Table 1).

The initial atomic model for protein residues of *Mtb* RNAP-Lpm was built by manual rigid-body fitting of RNAP β’, RNAP β, RNAP α^I^, RNAP α^II^, RNAP ω, and σ segments from the crystal structure of *Mtb* RPo (PDB 5UH5^15^; protein residues only) into the cryo-EM density map in Coot^80^, followed by adjustment of backbone and sidechain conformations in Coot. For σR4 (residues 464-528), density was weak, suggesting high segmental flexibility; σR4 was fitted as a rigid-body segment and was not further adjusted. For the RNAP β’ N and C-termini (residues 1-2 and 1282-1316), the central part of the RNAP β’ trigger loop (residues 1014-1022), the RNAP β N and C-termini (residues 1-27 and 1145-1172), the RNAP α^I^ N-terminus and C-terminal domain (residues 1-2 and 227-347), the RNAP α^II^ N-terminus and C-terminal domain (residues 1-2 and 233-347), the RNAP ω N-terminus (residues 1-27), σR1.1 (residues 1-224), and a loop and short extended segment of the σR3-σR4 linker (residues 426-433 and 443-445), density was absent, suggesting very high segmental flexibility; these segments were not fitted. The initial atomic model for Lpm atoms of *Mtb* RNAP-Lpm was built by manual rigid-body fitting of a crystal structure of Lpm (CCDC 114782^81^) into the cryo-EM density map using Coot^80^, followed by torsion-angle adjustments using Coot. Iterative cycles were performed of real-space model building in Coot^80^ followed by reciprocal-space fitting to structure-factor amplitudes and phases calculated from the cryo-EM density map in Phenix^82^. The final atomic model, with map-to-model correlation of 0.83, was deposited in the Electron Microscopy Data Bank (EMDB) and Protein Data Bank (PDB) with accession codes EMDB 4230 and PDB 6FBV (Extended Data Table 1).

### Crystal structure determination *(T. thermophilus* RNAP core enzyme): sample preparation

Robotic crystallization trials were performed for *T. thermophilus* RNAP core enzyme using a Gryphon liquid handling system (Art Robbins Instruments, Inc.), commercial screening solutions (Emerald Biosystems, Hampton Research, and Qiagen), and the sitting-drop vapor-diffusion technique (drop: 0.2 μl RNAP plus 0.2 μl screening solution; reservoir: 60 μl screening solution; 22°C). 900 conditions were screened. Rod-like crystals appeared under the identified crystallization conditions [0.1 M Hepes-NaOH, pH 7.5, 20 mM MgCl_2_, and 22% poly(acrylic acid sodium salt), 5,100 Da (Hampton Research); 22°C] within two weeks. Crystals were transferred from the sitting drop to a reservoir solution containing 20% (v/v) ethylene glycol (Sigma-Aldrich) and flash-cooled by immersing in liquid nitrogen.

### Crystal structure determination *(T. thermophilus* core enzyme): data collection and data reduction

Diffraction data and selenium single-wavelength anomalous dispersion data were collected from cryo-cooled crystals at Argonne Photon Source beamline 19ID-D. Data were processed using HKL2000^83^. The resolution cut-off criteria were I/σ > 1.1 and Rm_erge_ < 1.

The structure of T. *thermophilus* RNAP core was solved by molecular replacement with Molrep^84^ using PDB 4GZY^85^ as the search model. One RNAP molecule was present in the asymmetric unit. Early-stage refinement included rigid-body refinement of RNAP, followed by rigid-body refinement of RNAP subunits, followed by rigid-body refinement of 44 RNAP domains (methods as described^36^).

Cycles of iterative model building with Coot^80^ and refinement with Phenix^82^ were performed. Improvement of the coordinate model resulted from improvement of phasing. The final model was generated by XYZ-coordinate refinement with secondary-structure restraints in Phenix, followed by group B-factor and individual B-factor refinement in Phenix. The final model, refined to Rwork and Rfree of 0.22 and 0.27, respectively, was deposited in the PDB with accession code PDB 6ASG (Extended Data Table 2).

### Single-molecule FRET: sample preparation

Observation wells were prepared essentially as described^86–87^ (Extended Data Fig. 8b):

Borosilicate glass coverslips (#1.5; Menzel Gläzer) were incubated in 40 ml acetone 5 min at 22°C, incubated in 40 ml 2% (v/v) Vectabond aminosilane reagent (Vector Labs) in acetone 5 min at 22°C, washed with 100 ml water at 22°C, dried under nitrogen, and bonded to CultureWell 6 mm silicone gaskets (GBL103280; Grace Bio-labs), yielding 30 μl wells containing aminosilane-functionalized glass floors. Aliquots (20 μl) of 30 mM methoxy-PEG succinimidyl valerate (mPEG-SVA, MW 5,000; Laysan Bio) and 0.75 mM biotinyl-PEG succinimidyl valerate (Biotin-PEG-SVA, MW 5,000; Laysan Bio) in 50 mM MOPS-NaOH, pH 7.5, were added to wells and incubated 90 min at 22°C, supernatants were removed, and wells were washed with 5x200 μl PBS (0.01 M sodium phosphate, pH. 7.4, 137 mM NaCl, and 2.7 mM KC), yielding wells with biotin-PEG/mPEG-functionalized borosilicate glass floors. Aliquots (30 μl) of 10 μM Neutravidin (Sigma Aldrich) in 0.5xPBS then were added to wells and incubated 10 min at 22°C, supernatants were removed, and wells were washed with 3x100 μl PBS, yielding wells with Neutravidin-biotin-PEG/PEG-functionalized glass floors. Aliquots (30 μl) of 40 nM biotinylated anti-hexahistidine monoclonal antibody (Penta-His Biotin Conjugate; Qiagen) in buffer KG7 (40 mM Hepes-NaOH, pH 7.0, 100 mM potassium glutamate, 10 mM MgCl_2_, 1 mM dithiothreitol, 100 μg/ml bovine serum albumin, and 5% glycerol) then were added to wells and incubated 10 min at 22°C, supernatants were removed, and wells were washed with 3x100 μl KG7, yielding wells with (biotinylated anti-hexahistidine monoclonal antibody)-Neutravidin-biotin-PEG/mPEG-functionalized glass floors.

For experiments in Fig. 3c-e and Extended Data Fig. 8c-d, fluorescent-probe-labelled, hexahistidine-tagged *E. coli* RNAP σ^70^ holoenzyme was immobilized in observation wells containing (biotinylated anti-hexahistidine monoclonal antibody)-Neutravidin-biotin-PEG/mPEG-functionalized glass floors (Extended Data Fig. 8b) as follows: Aliquots (30 μl) of 0.1 nM fluorescent-probe-labelled. hexahistidine-tagged *E. coli* RNAP σ^70^ holoenzyme and 0 or 20 μM Lpm in KG7 (prepared by pre-equilibrating 50 nM fluorescent-probe-labelled. hexahistidine-tagged *E. coli* RNAP σ^70^ holoenzyme and 0 or 20 μM Lpm in KG7 10 min at 37°C, and then diluting 1:500 with 0 or 20 uM Lpm in KG7 at 37°C) were added and incubated 2-4 min at 22°C, supernatants were removed, wells were washed with 2x30 μl KG7, and 30 μl KG7 containing 2 mM Trolox (Sigma-Aldrich) and an oxygen scavenging system [12.5 μM glucose oxidase (Sigma-Aldrich), 16 nM catalase (C30; Sigma-Aldrich), and 8 mM D-glucose] at 22°C was added. Immobilization densities typically were ~30 molecules per 10 μm x 12 μm field of view. Immobilization specificities typically were >98% (assessed in control experiments omitting biotinylated anti-hexahistidine monoclonal antibody).

For experiments in Fig. 3f, fluorescent-probe-labelled, hexahistidine-tagged *E. coli* RNAP σ^70^ holoenzyme was immobilized in observation wells containing (biotinylated anti-hexahistidine monoclonal antibody)-Neutravidin-biotin-PEG/mPEG-functionalized glass floors in the absence of Lpm as described above, and 3 μl 200 μM Lpm in the same buffer was added during data acquisition (Lpm final concentration = 20 μM).

### Single-molecule FRET: data collection and data analysis

FRET experiments were performed on an objective-type total-internal-reflection (TIRF) microscope^86–88^. Light from a green laser (532 nm; Samba; Cobolt) and a red laser (635 nm; CUBE 635-30E; Coherent) was combined using a dichroic mirror, coupled into a fiber-optic cable, focused onto the rear focal plane of a 100x oil-immersion objective (numerical aperture 1.4; Olympus), and displaced off the optical axis such that the incident angle at the oil-glass interface of a stage-mounted observation chamber was greater than the critical angle, thereby creating an exponentially decaying evanescent wave. Alternating-laser excitation (ALEX^89^) was implemented by directly modulating the two lasers using an acousto-optical modulator (1205C; Isomet). Fluorescence emission was collected from the objective, separated from excitation light by a dichroic mirror (545 nm/650 nm; Semrock) and emission filters (545 nm LP, Chroma; and 633/25 nm notch filter, Semrock), focussed on a slit to crop the image, and then spectrally separated using a dichroic mirror (630 nm DLRP; Omega) into donor and emission channels focused side-by-side onto an electron-multiplying charge-coupled device camera (EMCCD; iXon 897; Andor Technology). A motorized x/y-scanning stage with continuous reflective-interface feedback focus (MS-2000; ASI) was used to control the sample position relative to the objective.

For experiments in Fig. 3c-e and Extended Data Fig. 8c-d, laser powers were 3.5 mW (532 nm laser) and 0.7 mW (635 nm laser), and data were collected for 20 s using a frame rate of 1 frame per 20 ms. For experiments in Fig. 3f, laser powers were 0.5 mW (532 nm laser) and 0.15 mW (635 nm laser), and data were collected for 50 s using a frame rate of 200 ms.

Fluorescence emission intensities in donor (green) and acceptor (red) emission channels were detected using the peak-finding algorithm of the MATLAB (MathWorks) software package Twotone-ALEX^88^, as described^88^. Peaks detected in both emission channels (i.e., peaks for molecules containing both donor and acceptor probes) and meeting ellipticity and distance-to-nearest-neighbor thresholds (i.e., ellipticity ≤ 0.6 and distance-to-nearest-neighbor ≥ 6 pixels) were fitted with two-dimensional Gaussian functions to extract background-corrected intensity-vs.-time trajectories for donor emission intensity upon donor excitation (I_DA_), acceptor emission intensity upon donor excitation (I_DD_), and acceptor emission intensity upon acceptor excitation (I_AA_) (Extended Data Fig. 8c, top), as described^88^. Intensity-vs.-time trajectories were curated to exclude trajectories exhibiting I_DD_ < 300 or >2,000 counts or I_AA_ < 200 or >2,000 counts, trajectories exhibiting multiple-step donor or acceptor photobleaching, trajectories exhibiting donor or acceptor photobleaching in frames 1-50, and trajectories exhibiting donor or acceptor photoblinking, and to exclude portions of traces following donor or acceptor photobleaching.

Intensity-vs.-time trajectories were used to calculate trajectories of apparent donor-acceptor FRET efficiency (E*) and donor-acceptor stoichiometry (S) (Extended Data Fig. 8c, bottom), as described^89–90^:

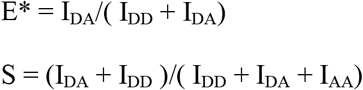

E*-vs.-S plots were prepared, S values were used to distinguish species containing only donor, only acceptor, and both donor and acceptor, and E* histograms were prepared for species containing both donor and acceptor, as described^89–90^.

E*-vs. time trajectories that on visual inspection exhibited transitions between distinct E* states (dynamic E*-vs.-time trajectories; ~34%; N = 207) were analyzed globally to identify E* states by use of Hidden Markov Modelling (HMM) with an empirical Bayesian as implemented in the MATLAB (MathWorks) software package ebFRET^91^, essentially as described^86–87, 91^. E*-vs.-time trajectories were fitted to HMM models with two, three, four, five, or six distinct E* states; the mean scoring parameter L (lower bound per trajectory) was extracted for each model. and the model with three distinct E* states was found to best describe the data (Fig. 3c, right; L = 536, 538, 537,537 and 537 for models with two, three, four, five, and six states). E* values from the three-state HMM model were extracted, plotted using Origin (OriginLab), and fitted to Gaussian distributions using Origin (Fig 3c, left, colored curves). The resulting histograms provide equilibrium population distributions of E* states and, for each E* state, define mean E* (Fig 3c, left, colored curves and inset).

Dwell times for E* states were extracted from E*-vs.-time trajectories exhibiting >3 transitions between E* states and were plotted as histograms in Origin (Extended Data Fig. 8d). The resulting dwell-time histograms were fit with single-exponential functions, and mean dwell times were extracted (Fig. 3e; Extended Data Fig. 8d).

E* values were corrected, and accurate donor-acceptor efficiencies (E) and donor-acceptor distances (R) were calculated, as follows^90^.

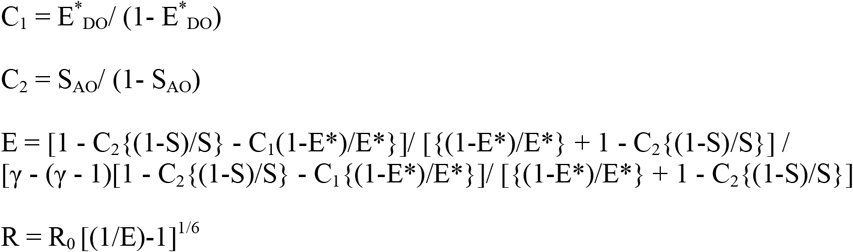

where, C_1_ is “cross-talk” from leakage of donor emission into the acceptor-emission channel, C_2_ is “cross-talk” from direct excitation of the acceptor by the green laser, γ is the detection factor (1 in this work; determined as γ = ΔI_AA_/ΔI_DD_, where ΔI_AA_ and ΔI_DD_ are changes in I_AA_ and I_DD_ upon acceptor photobleaching^92^), E*_DO_ is E* of the donor-only subpopulation, S_AO_ is S of the acceptor-only subpopulation, and R_0_ is the Főrster parameter [60.1 Å in this work; calculated as: R_0_= 9780(n^−4^κ^2^Q_D_J)^1/6^ Å, where n is the refractive index of the medium, κ^2^ is the orientation factor relating donor emission dipole and acceptor excitation dipole (approximated as 2/3, noting that all mean E values are <0.5^93^), Q_D_ is the quantum yield of the donor in the absence of acceptor, and J is the overlap integral of donor emission spectrum and acceptor excitation spectrum].

RNAP clamp angles were determined from E values using the plot of RNAP clamp angle vs. E in Fig. 1 of ref. 14.

### Data analysis

Data for RNAP-inhibitory activities, resistance, and cross-resistance are means of at least two technical replicates

### Data availability

The cryo-EM map and atomic coordinates for the cryo-EM structure of *Mtb* RNAP-Lpm have been deposited in the EMDB and PDB with accession codes EMDB 4230 and PDB 6FBV. Atomic coordinates and structure factors for the crystal structure of *T. thermophilus* RNAP core enzyme have been deposited in the PDB with accession code PDB 6ASG.

## EXTENDED DATA FIGURE LEGENDS

**Extended Data Figure 1.**
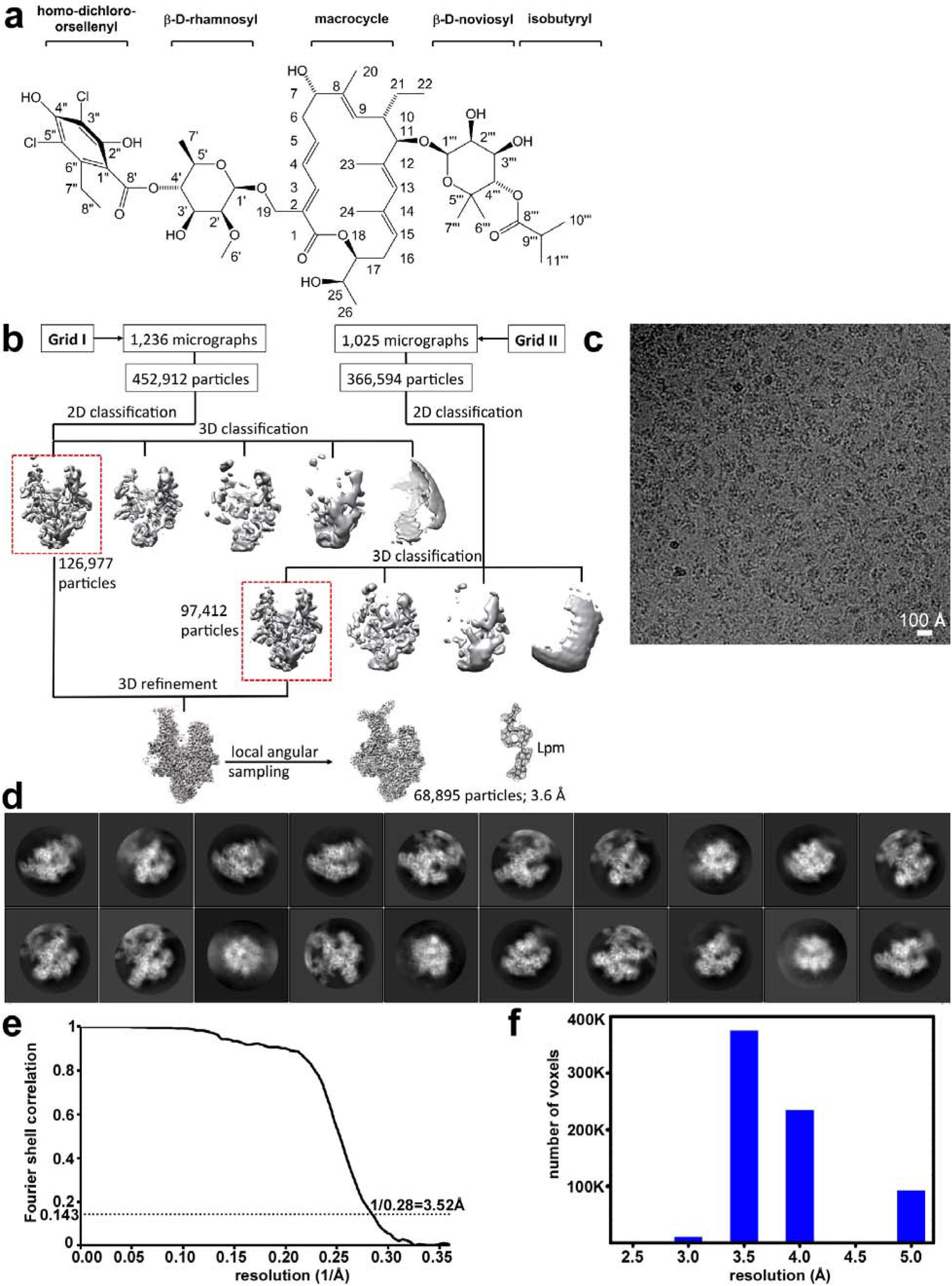
Structure of *Mtb* RNAP-Lpm: structure determination. **a**, Structure of lipiarmycin A3^94^ (Lpm). Atom numbering as described^94^. **b**, Summary of structure determination procedures. **c**, Representative electron micrograph. **d**, Representative two-dimensional classes. **e**, Fourier-shell-correlation (FSC) analysis. **f**, Resolution distribution.

**Extended Data Figure 2.**
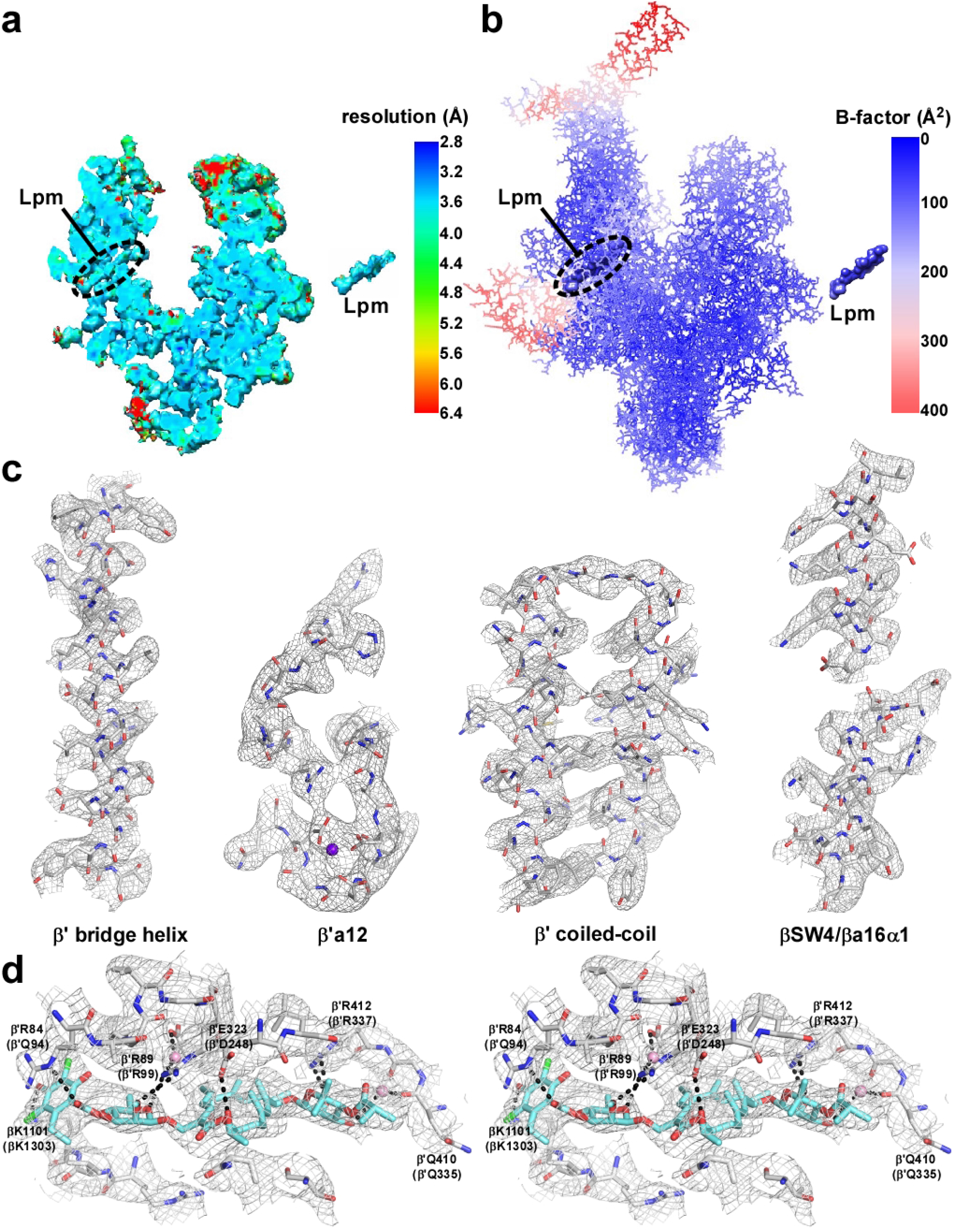
Structure of *Mtb* RNAP-Lpm: structure quality. **a**, Slice through density map of *Mtb* RNAP-Lpm at level of Lpm (left) and portion of slice comprising Lpm (right), colored by local resolution. View orientation as in Fig. 1. **b**, Structure of *Mtb* RNAP-Lpm colored by B-factor. View orientation as in a. **c**, Density maps for structural elements of central portion of *Mtb* RNAP-Lpm [two structural elements from RNAP active-center region (first and second panels) and two structural elements from RNAP clamp and RNAP switch region (third and fourth panels)]. Gray mesh, density map; gray, red, and green sticks, carbon, oxygen and nitrogen atoms; violet sphere, RNAP active-center catalytic Mg^2+^. **d**, Density map for Lpm binding site (stereodiagram). Cyan and green sticks, Lpm carbon and chloride atoms; black dashed lines, H-bonds. Other colors as in **c**.

**Extended Data Figure 3.**
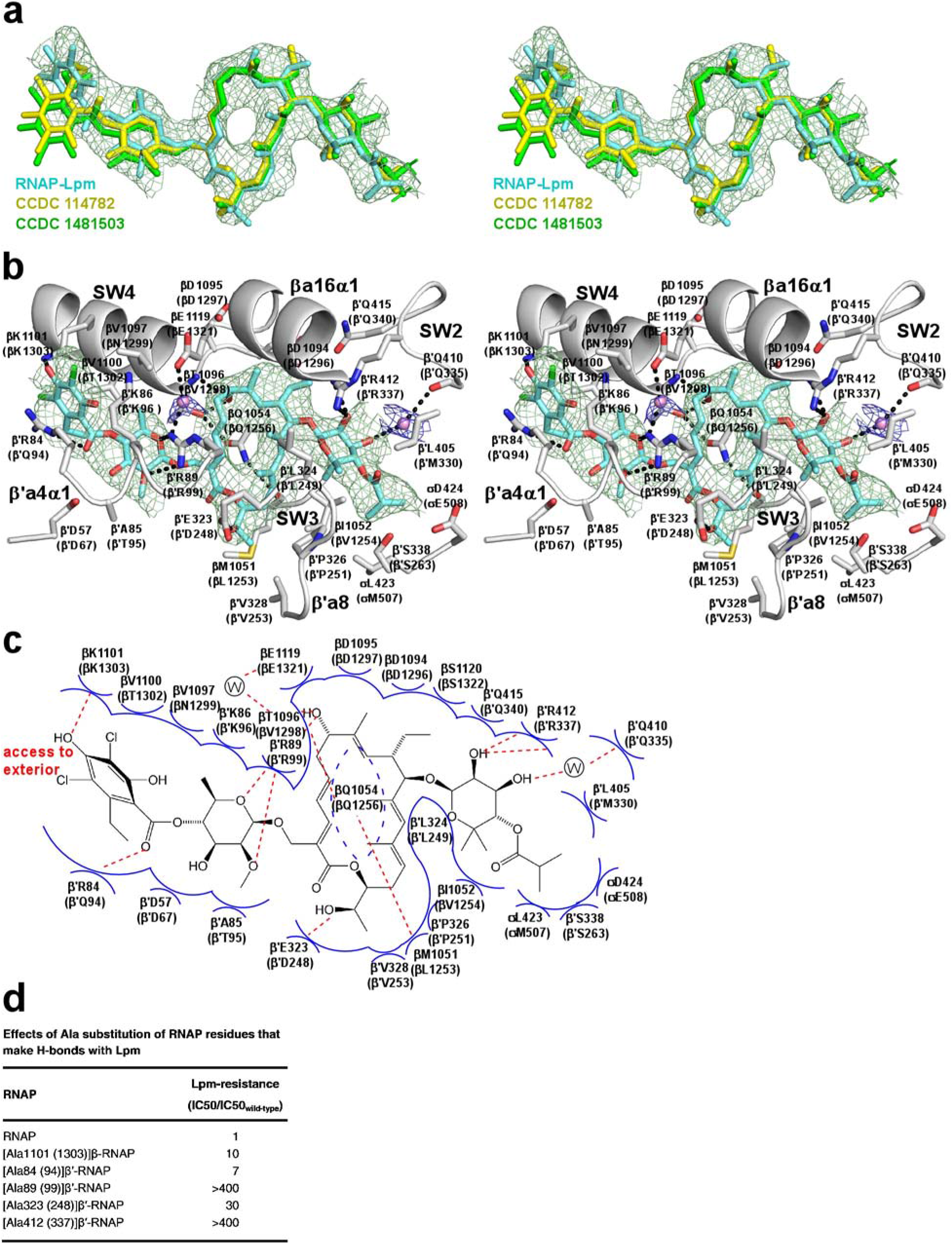
Structure of Mtb RNAP-Lpm: RNAP-Lpm interactions. **a**, Stereoview of density and atomic model for Lpm in *Mtb* RNAP-Lpm (green mesh and cyan sticks), superimposed on atomic models for Lpm from crystal structures of Lpm alone (yellow and green sticks; CCDC 114782 and CCDC 1481503). **b**, Stereoview of inferred RNAP-Lpm interactions. Image extends information in Fig. 1d by including density and fit for interfacial water molecules (blue mesh and pink spheres) and annotations for RNAP structural elements (SW2, SW3, SW4, βa16α1, β’a4α1, and β’a8). **c**, Summary of inferred RNAP-Lpm interactions. Image extends information in Fig. 1e by including interfacial water molecules (“W”) and indicating solvent exposure of the Lpm C4” hydroxyl and C5” chlorine. **d**, Effects of alanine substitution of RNAP residues inferred to make H-bonds with Lpm. Residues in **b-d** are numbered as in *Mtb* RNAP and, in parentheses, as in *E. coli* RNAP.

**Extended Data Figure 4.**
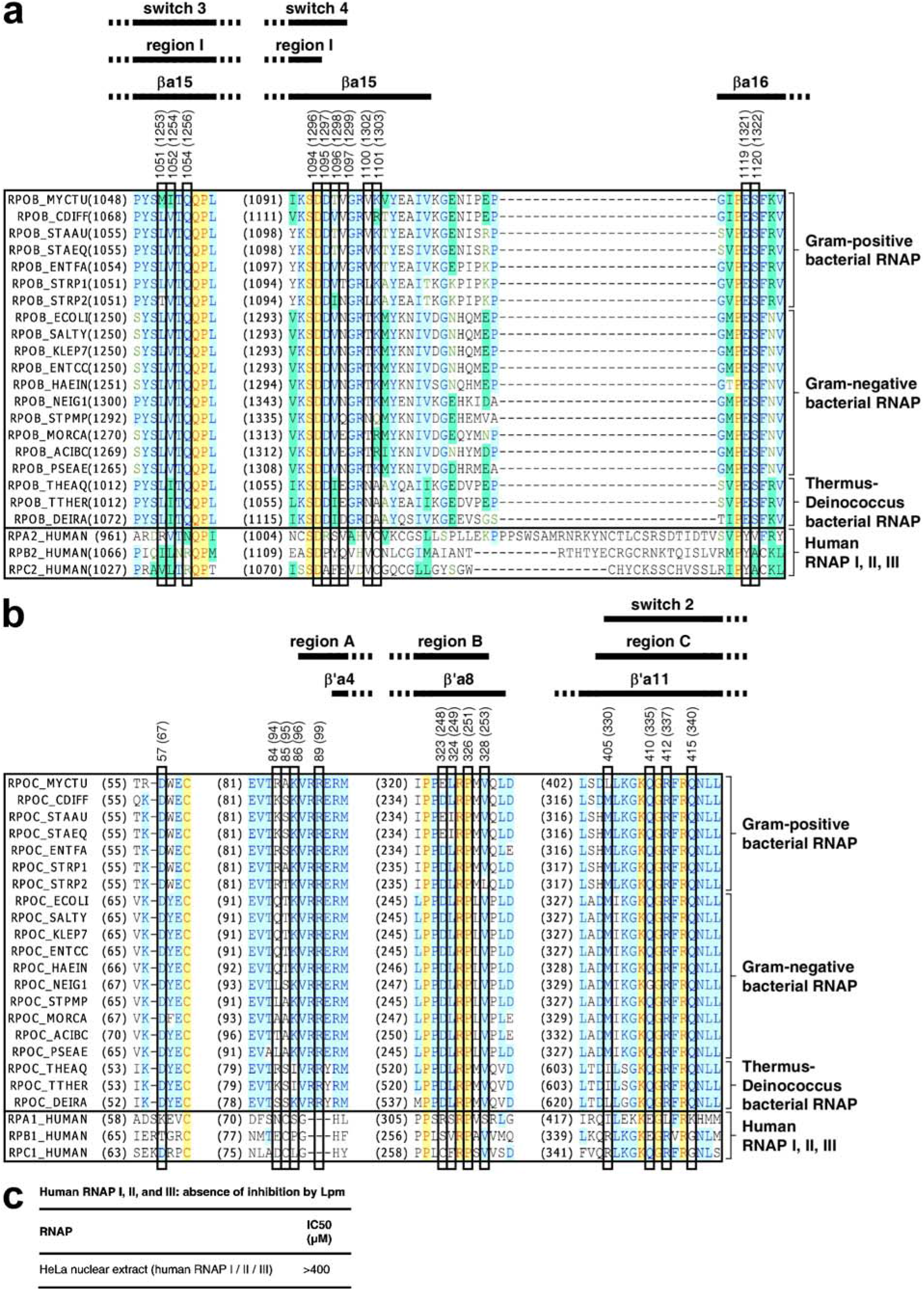
Sequence alignments and species selectivity of inhibition. **a-b**, Locations of residues that contact Lpm in sequences of RNAP β subunit (**a**) and RNAP β’ subunit (**b**). Sequence alignments for the β and β’ subunits of bacterial RNAP (top 20 sequences in each panel) and corresponding subunits of human RNAP I, RNAP II, and RNAP III (bottom three sequences in each panel), showing locations of RNAP residues that contact Lpm (black rectangles; identities from Fig. 1d-e and Extended Data Fig. 3b-c), RNAP switch-region SW2-SW4 (first row of black bars; boundaries as described^17^), and RNAP conserved regions and conserved structural elements (second and third rows of black bars; boundaries as described^95–97^). Species are as follows: *Mycobacterium tuberculosis* (MYCTU), *Clostridium difficile* (CDIFF), *Staphylococcus aureus* (STAAU), *Staphylococcus epidermidis* (STAEQ), *Enterococcus faecalis* (ENTFA), *Streptococcus pyogenes* (STRP1), *Streptococcus pneumoniae* (STRP2), *Escherichia coli* (ECOLI), *Salmonella typhimurium* (SALTY), *Klebsiella pneumoniae* (KLEP7), *Enterococcus cloacae* (ENTCC), *Vibrio cholerae* (VIBCH), *Haemophilus influenzae* (HAEIN), *Neisseria gonorrhoeae* (NEIG1), *Stenotrophomonas maltophilia* (STPMP), *Moraxella catarrhalis* (MORCA), *Acinetobacter baumannii* (ACIBC), *Pseudomonas aeruginosa* (PSEAE), *Thermus thermophilus* (THETH), *Thermus aquaticus* (THEAQ), *Deinococcus radiodurans* (DEIRA), and *Homo sapiens* (HUMAN). **c**, Absence of inhibition of human RNAP I, RNAP II, and RNAP III by Lpm.

**Extended Data Figure 5.**
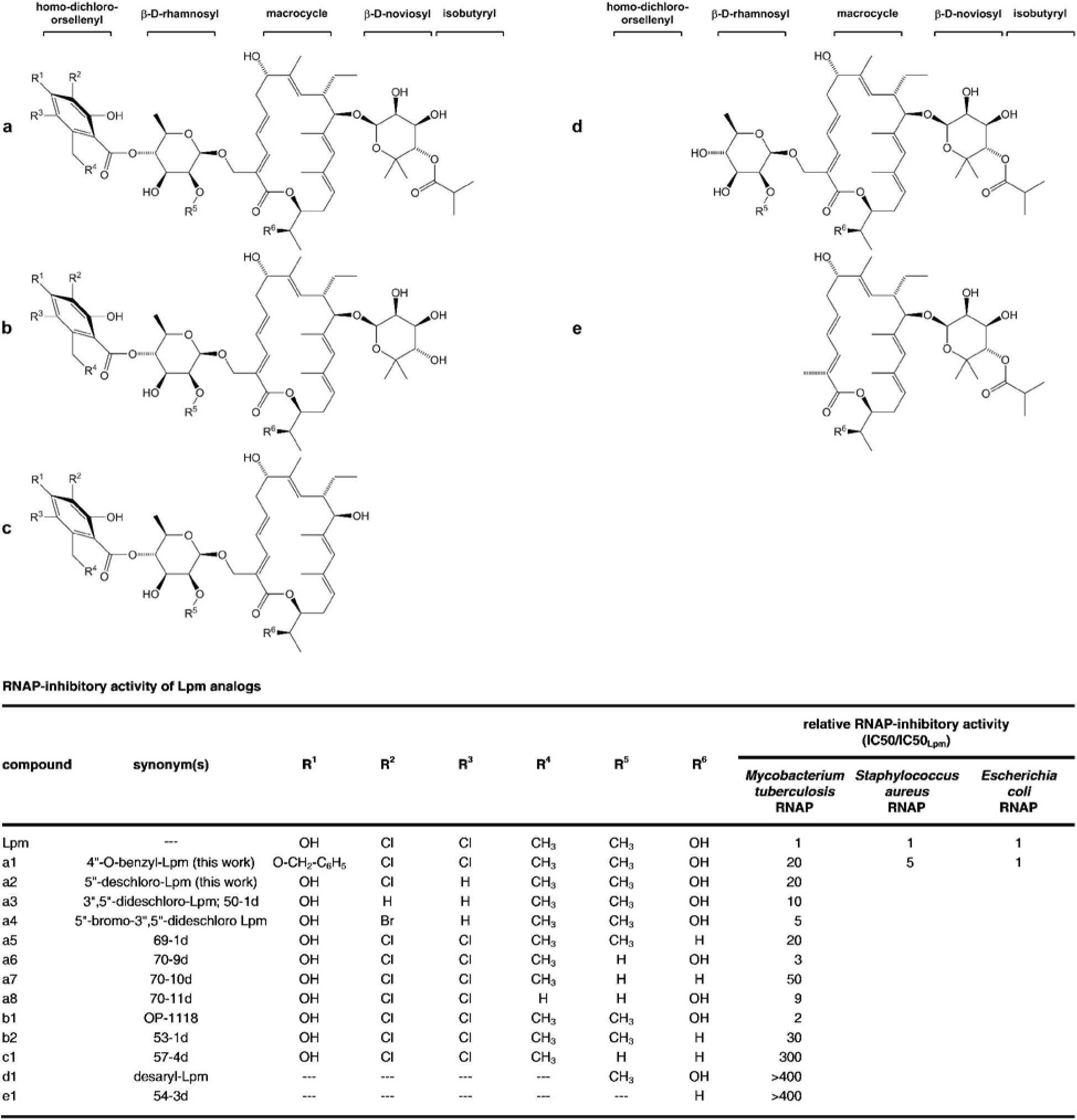
Structure-activity relationships for Lpm analogs. **a**, Structures of Lpm analogs. Compound “a”1 is Lpm.^94^ Compounds “a3”-”c1” and “e1” are previously known Lpm analogs^18–21^. Compounds “a1”-”a2” and ““d1” are new Lpm analogs (see Methods). **b**, RNAP-inhibitory activities of Lpm analogs.

**Extended Data Figure 6.**
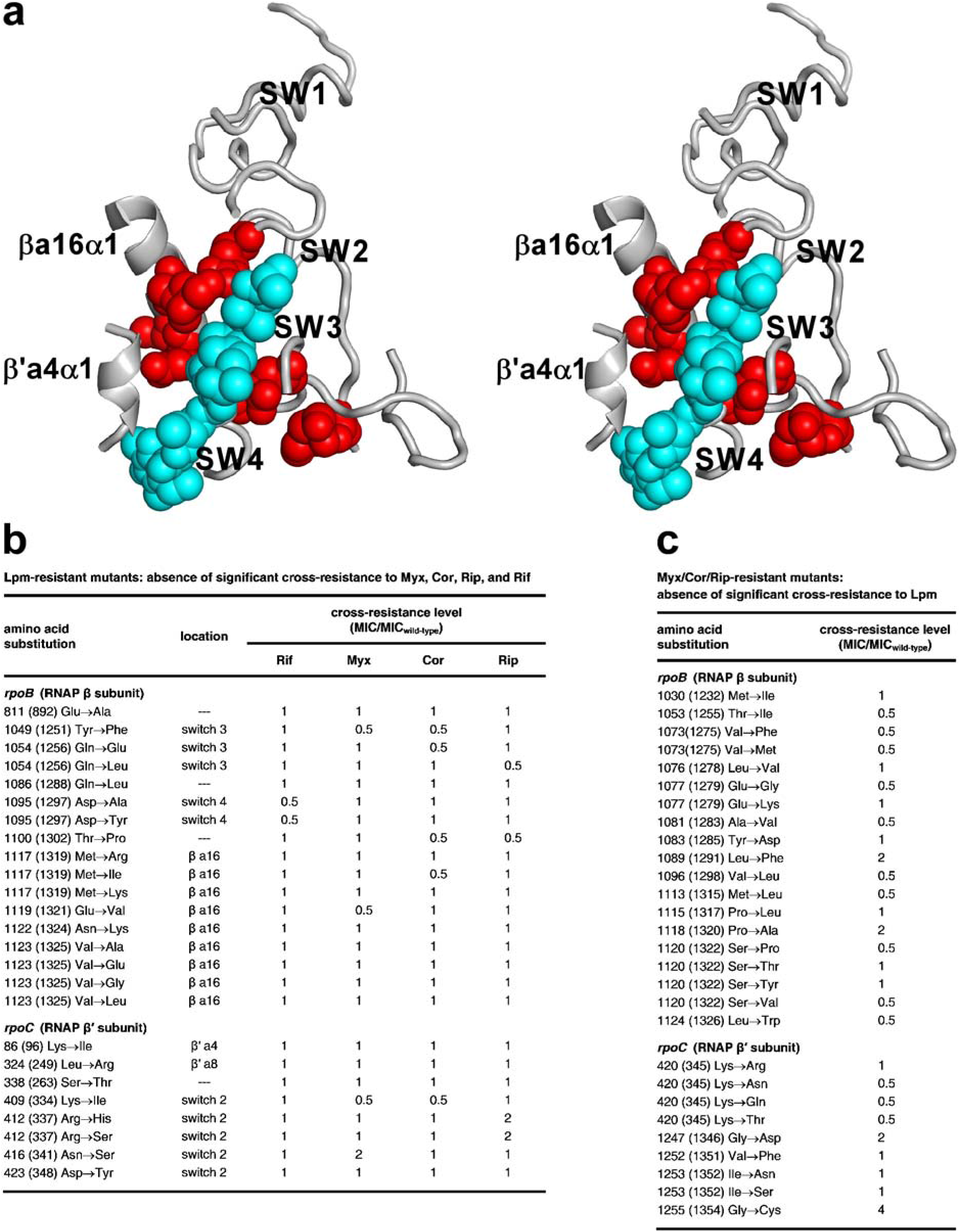
Lpm-resistance and cross-resistance. **a**, Stereodiagram of RNAP-Lpm interactions, showing RNAP structural elements SW2, SW3, SW4, βa16α1, and β’a4α1, and showing amino acids at which substitutions conferring high-level (≥4-fold) Lpm-resistance are obtained (sequences from Fig. 2c). Gray ribbons, RNAP; cyan, Lpm; red, resistance sites. **b**, Comprehensive analysis of cross-resistance of Lpm-resistant mutants (sequences from Fig. 2c) to Rif, Myx, Cor, and Rip. **c**, Comprehensive analysis of cross-resistance of Myx-, Cor-, and Rip-resistant mutants to Lpm. [Substitution of RNAP β 1123 (1325) to Leu previously was reported to confer resistance to Myx, Cor, and Rip, but re-analysis indicates no resistance to Myx, Cor, and Rip.] Residues in **b-c** are numbered as in *Mtb* RNAP and, in parentheses, as in *E. coli* RNAP.

**Extended Data Figure 7.**
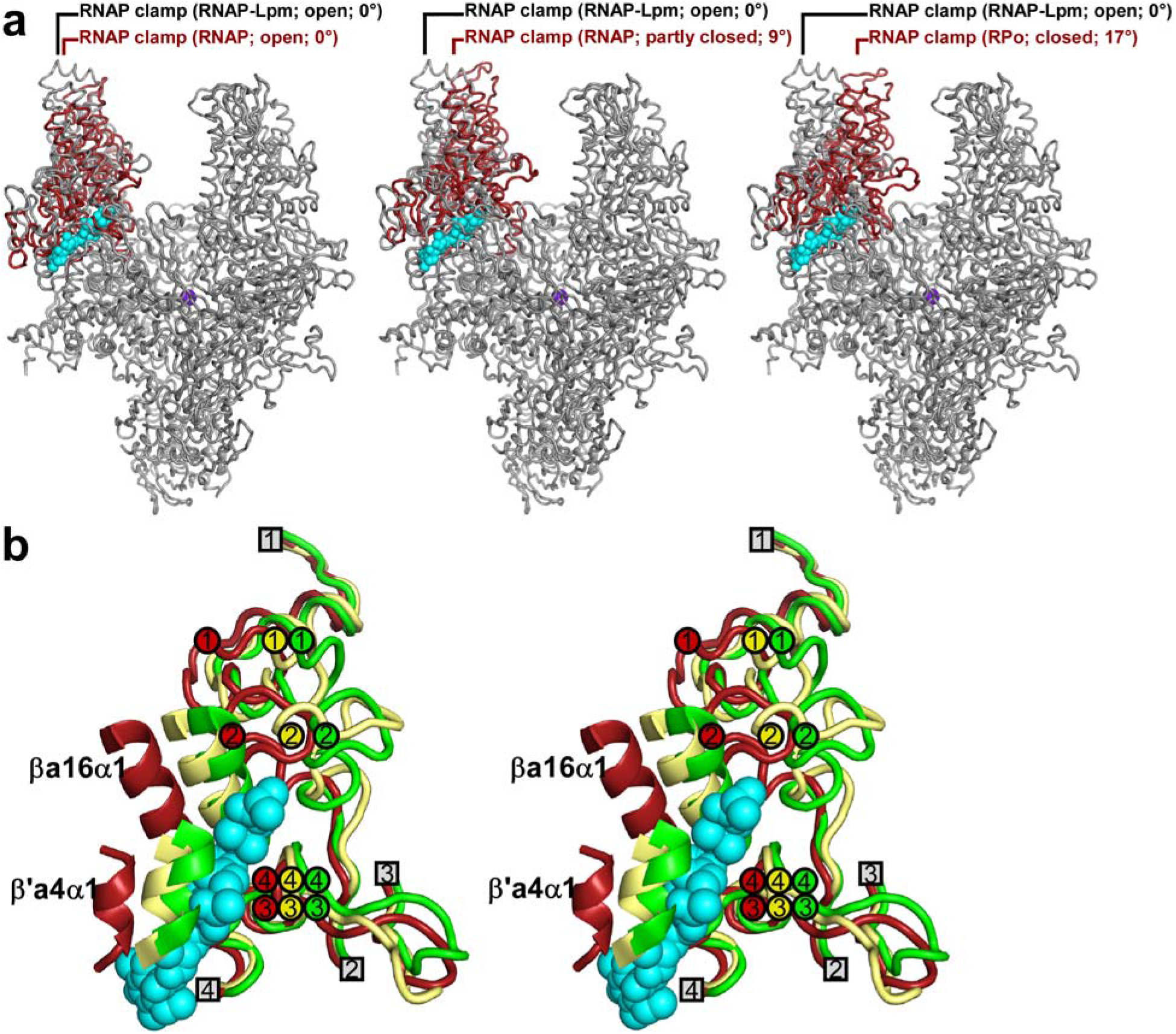
Effect of Lpm on RNAP clamp conformation: cryo-EM. **a**, Comparisons of RNAP clamp conformation in *Mtb* RNAP-Lpm to RNAP clamp conformations in crystal structures of *Tth* RNAP with open clamp (left; Extended Data Table 2), *Tth* RNAP with partly closed clamp (center; PDB 5TMC), and *Tth* RPo with closed clamp (right; PDB 4G7H). Each image shows *Mtb* RNAP-Lpm (view orientation and colors as in Fig. 1–2) and, superimposed, the RNAP clamp of the comparator structure (red). **b**, Stereodiagram of Lpm binding site showing conformations of RNAP structural elements SW1, SW2, SW3, SW4, βa16α1, and β’a4α1, in open-clamp (red), partly-closed clamp (yellow), and closed-clamp (green) conformational states. RNAP switches. SW1, SW2, SW3, and SW4 are shown with ends that connect to the RNAP clamp as numbered circles, and ends that connect to the RNAP main mass as numbered squares. The open-clamp state is from the structure of *Mtb* RNAP-Lpm (Fig. 1c); the partly-closed-clamp and closed-clamp states are from PDB 5TMC and PDB 4G7H. Lpm is in cyan..

**Extended Data Figure 8.**
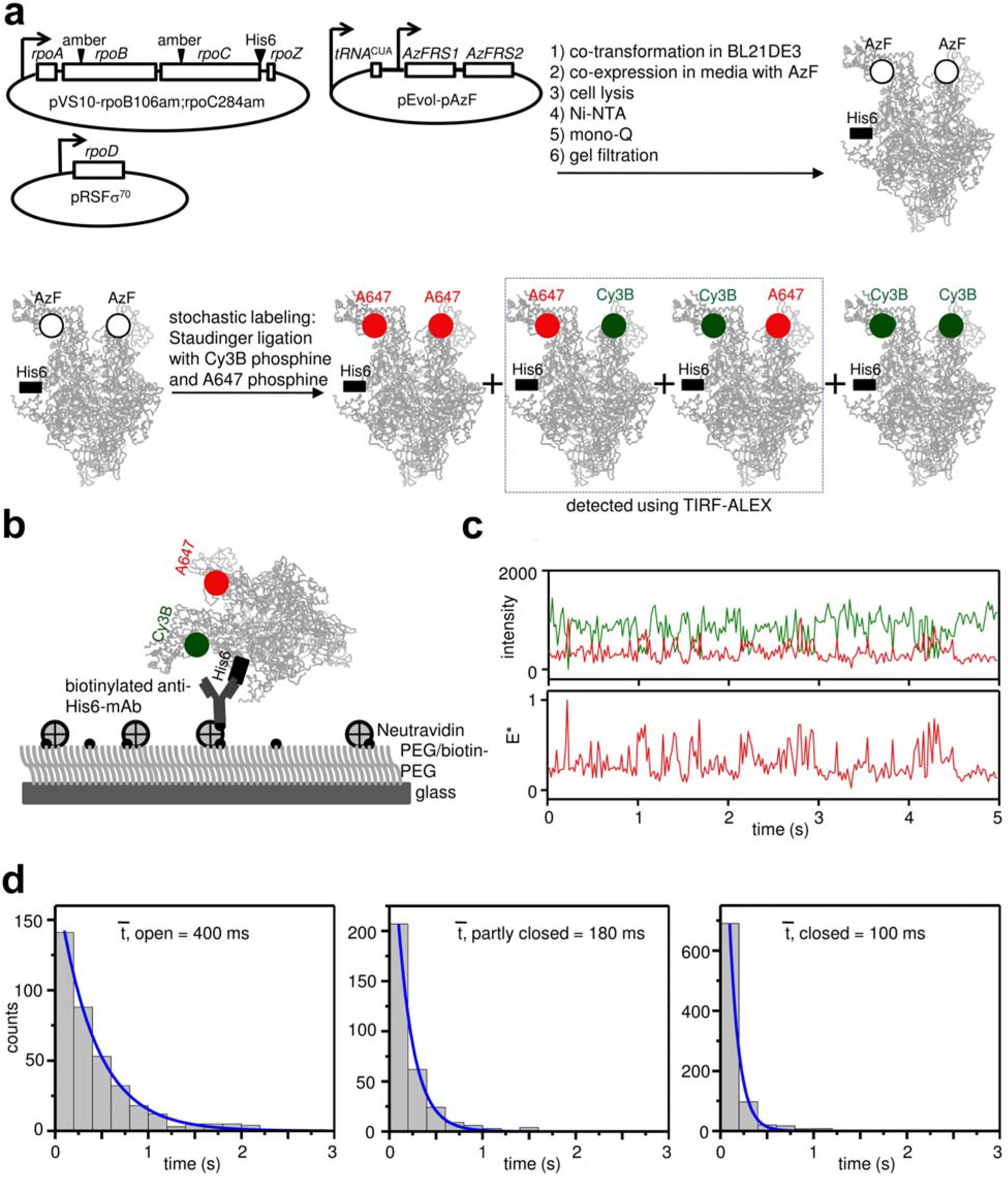
Effect of Lpm on RNAP clamp conformation: single-molecule FRET. **a**, Use of unnatural-amino-acid mutagenesis^14, 52^, stochastic Staudinger ligation with Cy3B-phosphine and Alexa647-phosphine^14, 53–55^, and total-internal-reflection fluorescence microscopy with alternating-laser excitation microscopy (TIRF-ALEX)^86–88^ to obtain FRET data for single molecules of RNAP having one fluorescent probe at the tips of the walls of the RNAP active-center cleft. Green, fluorescence donor probe Cy3B; red, fluorescent acceptor probe Alexa647; black square, hexahistidine tag. **b**, Surface-immobilization of fluorescent-probe-labelled RNAP for ALEX-TIRF. **c**, Representative time traces of donor emission intensity (green) and acceptor emission intensity (red) (top) and corresponding time trace of computed donor-acceptor FRET efficiency (bottom). **d**, Dwell-time distributions and extracted mean dwell times for HMM-assigned open-clamp (left), partly-closed-clamp (center), and closed-clamp (right) states.

**Extended Data Table 1.** Cryo-EM data-collection and data-analysis statistics

|  |  |  |
| --- | --- | --- |
| **subject** | *Mtb* RNAP-Lpm |  |
| **EMDB and PDB codes** | 6FBV and 4230 |  |
| **data collection** |  |  |
| number of grids used | 2 |  |
| grid type | lacey carbon |  |
| microscope/detector | Titan Krios/Gatan K2 |  |
| voltage (kV) | 300 kV |  |
| magnification | 130,000x |  |
| recording mode | Counting mode |  |
| dose rate (e-/s) | 8 |  |
| pixel size (Å/pix) | 1.061 |  |
| total dose (e-/Å<sup>2</sup>) | 72.05 |  |
| number of frames/movie | 50 |  |
| total exposure time (s) | 10 |  |
| defocus range (μm) | 1.0 - 2.0 |  |
| **data processing** | **grid I** | **grid II** |
| number of micrographs | 1,238 | 1,029 |
| number of particles picked | 452,912 | 366,594 |
| number of particles after 2D/3D | 126,977 | 97,412 |
| combined dataset |  |  |
| particles used for final map calculation | 68,895 |  |
| map resolution (FSC 0.143; Å) | 3.52 |  |
| map resolution (ResMap median; Å) | 3.5 |  |
| **structure/map fitting** |  |  |
| resolution (Å) | 3.5 |  |
| map sharpening B factor (Å<sup>2</sup>) | -109 |  |
| experimental map / model correlation | 0.83 |  |
| experimental map / 2Fo-Fc correlation | 0.98 |  |
| average B factor (Å<sup>2</sup>) | 95.3 |  |
| total number of atoms | 25,156 |  |
| clash score | 13 |  |
| Ramachandran plot |  |  |
| favored (%) | 94.5 |  |
| outliers (%) | 0.03 |  |
| root-mean-square (RMS) deviations |  |  |
| bond length (Å) | 0.006 |  |
| bond angle (°) | 1.027 |  |

\*Structure factors were from cryoEM map, and 2Fo-Fc map was from final model.

**Extended Data Table 2.** X-ray crystallography data-collection and data-analysis statistics*

|  |  |
| --- | --- |
| **subject** | *Tth* core |
| **PDB code** | 6ASG |
| data collection source | APS-19ID |
| **data collection** |  |
| space group | R3 |
| cell dimensions |  |
| a, b, c (Å) | 280.7,280.7,184.9 |
| $\alpha$, $\beta$, $\gamma$ (°) | 90.0, 90.0, 120.0 |
| resolution | 50.0-3.8 (4.0-3.8) |
| number of unique reflections | 61,395 |
| $R_{\text{merge}}$ | 0.245 (0.704) |
| $R_{\text{meas}}$ | 0.275 (0.830) |
| $R_{\text{pim}}$ | 0.124 (0.430) |
| highest resolution shell $CC_{1/2}$ | 0.418 |
| $I/\sigma(I)$ | 5.0 (1.2) |
| completeness (%) | 99.2 (92.3) |
| redundancy | 4.8 (3.2) |
| **refinement** |  |
| resolution (Å) | 42.5-3.8 |
| number of unique reflections | 50,511 |
| number of test reflections | 2,611 |
| $R_{\text{work}}/R_{\text{free}}$ | 0.22/0.27 (0.27/0.32) |
| number of atoms |  |
| protein | 23,551 |
| ligand/ion | 3 |
| root-Mean-Square(RMS) Deviations |  |
| bond length (Å) | 0.003 |
| bond angle (°) | 0.658 |
| MolProbity statistics |  |
| clash score | 15.2 |
| rotamer outliers (%) | 1.0 |
| Cβ outliers (%) | 0 |
| Ramachandran plot |  |
| Favored (%) | 94.1 |
| Outliers (%) | 0.4 |

\*Data for highest-resolution shells are in parentheses.

